# Novel Role of UHRF1 in DNA methylation-mediated repression of latent HIV-1

**DOI:** 10.1101/2021.06.15.448539

**Authors:** Roxane Verdikt, Sophie Bouchat, Alexander O. Pasternak, Lorena Nestola, Gilles Darcis, Véronique Avettand-Fenoel, Caroline Vanhulle, Amina Aït-Ammar, Maryam Bendoumou, Estelle Plant, Valentin Le Douce, Nadège Delacourt, Aurelija Cicilionytė, Coca Necsoi, Francis Corazza, Caroline Pereira Bittencourt Passaes, Christian Schwartz, Martin Bizet, François Fuks, Asier Sáez-Cirión, Christine Rouzioux, Stéphane De Wit, Ben Berkhout, Virginie Gautier, Olivier Rohr, Carine Van Lint

## Abstract

The multiplicity, heterogeneity and dynamic nature of HIV-1 latency mechanisms are reflected in the current lack of functional cure for HIV-1 and in the various reported *ex vivo* potencies of latency-reversing agents. Here, we investigated the molecular mechanisms underlying the potency of the DNA methylation inhibitor 5-aza-2’-deoxycytidine (5-AzadC) in HIV-1 latency reversal. Doing so, we uncovered specific demethylation CpG signatures induced by 5-AzadC in the HIV-1 promoter. By analyzing the binding modalities to these CpG, we revealed the recruitment of the epigenetic integrator UHRF1 to the HIV-1 promoter. We further demonstrated the role of UHRF1 in DNA methylation-mediated silencing of the latent HIV-1 promoter. As a proof-of-concept to this molecular characterization, we showed that pharmacological downregulation of UHRF1 in *ex vivo* HIV^+^ patient cell cultures resulted in potent reactivation of latent HIV-1. Together, we identify UHRF1 as a novel actor in HIV-1 gene silencing and highlight that it constitutes a new molecular target for HIV-1 curative strategies.

## INTRODUCTION

Combination antiretroviral therapy (cART) is the only therapeutic option available for HIV- 1infected individuals. If cART is efficient in suppressing viral replication and in prolonging the lifespan of infected individuals, the persistence of transcriptionally-silent proviruses, particularly in latently-infected resting memory CD4^+^ T cells, still prevents HIV-1 eradication (1–3). As such, much effort has been put in understanding the multiple molecular factors involved in viral latency to develop new anti-HIV therapeutic strategies. One such strategy relies on the use of latency-reversing agents (LRAs) that target repressors of HIV-1 gene expression, thereby inducing a controlled activation of latent reservoirs (4, 5).

The multifactorial process of HIV-1 silencing during latency is controlled in part by the viral transactivator Tat and by cellular transcription factors (TFs) binding sites (TFBS) present in the viral promoter, located in the 5’ long terminal repeat (5’LTR)(6). In addition, epigenetic processes controlling the chromatin architecture of latent HIV-1 proviruses play key roles in viral transcriptional silencing (7, 8). Two CpG islands (CGIs) present in the 5’LTR region and surrounding the transcription start site have been reported to be hypermethylated in latently-infected model T-cell lines (Fig. 1A), thus participating in the 5’LTR heterochromatinization during latency (9–11). Methylation of the HIV-1 promoter in patient cells has been reported in some studies (9, 10, 12) but other reports denied the implication of 5’LTR methylation *ex vivo* (13–15). To explain these contradictory results, recent studies proposed that patients clinical characteristics, such as duration of the infection (12) and duration of the antiretroviral treatment (16, 17), might influence the accumulation of DNA methylation in the 5’LTR. In agreement with these heterogeneous profiles of DNA methylation on the HIV-1 promoter *ex vivo*, we have previously shown that latency reversal with the DNA methylation inhibitor 5-aza-2’-deoxycytidine (5-AzadC or decitabine) is associated with patient-specific qualitative and quantitative variations in HIV-1 reactivation from latency (18).

**Figure 1:**
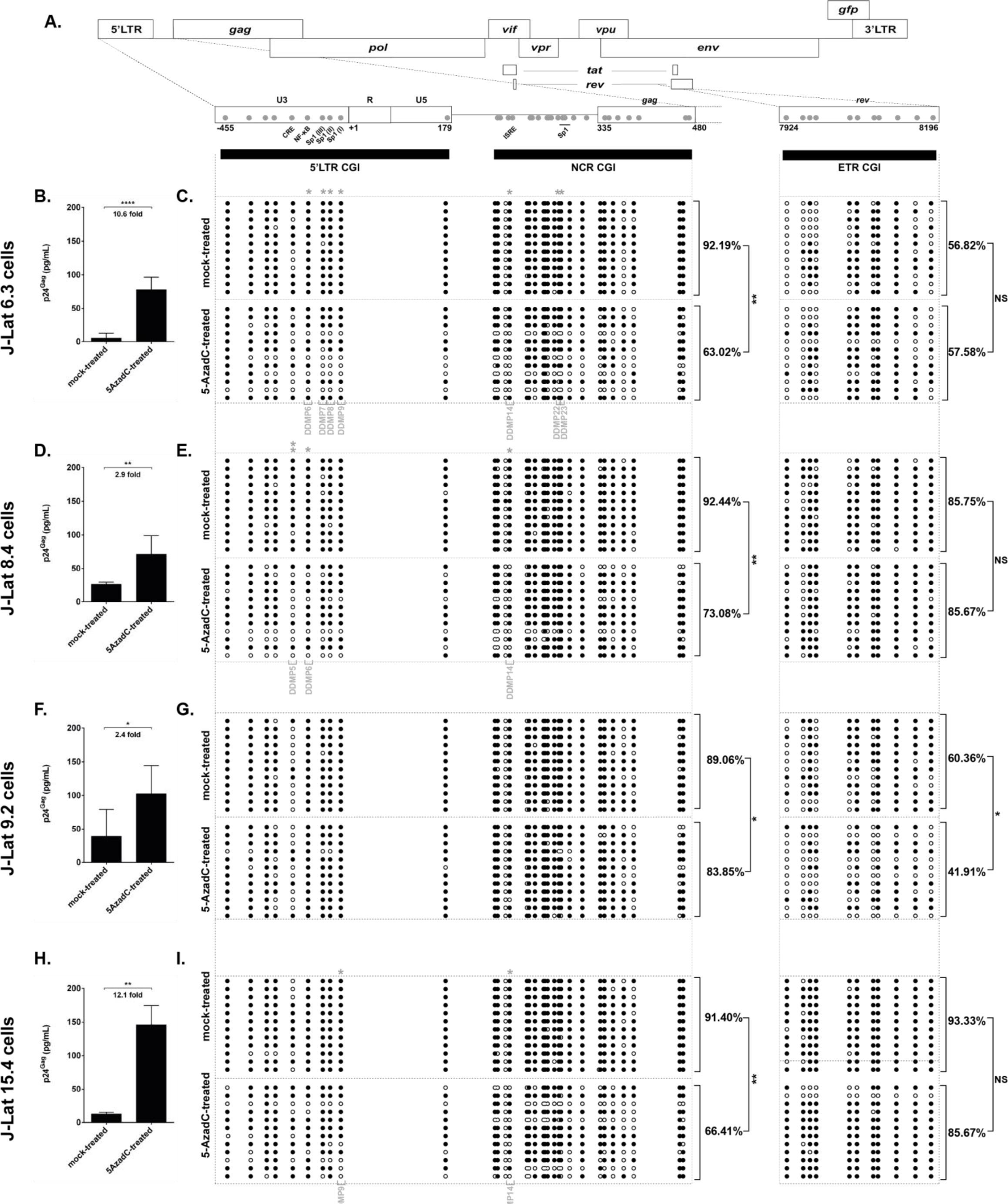
5-AzadC-induced reactivation of HIV-1 gene expression from latency is associated with 5’LTR CGIs demethylation. **(A)** Schematic presentation of the three CpG islands studied along the HIV-1 provirus, in the HIV-1 promoter (5’LTR and NCR CGIs) and in *rev* (ETR CGI). The reactivation of HIV-1 production following 72h treatment with 400nM of 5-AzadC, quantified by ELISA on p24^Gag^ capsid protein in culture supernatants, and the DNA methylation profile, established by sodium bisulfite sequencing for the three CGIs, are respectively presented for the J-Lat 6.3 cells **(B and C)**, the J-Lat 8.4 cells **(D and E)**, the J-Lat 9.2 cells **(F and G)** and the J-Lat 15.4 cells **(H and I)**. ELISA results are representative of the means ± SD of three independent 5-AzadC treatments. Folds reactivation are indicated. Unmethylated and methylated CpG dinucleotides are respectively represented with open and closed circles, where each line corresponds to individual sequenced molecules. The global methylation level presented correspond to mean percentages of methylated CpGs for the twelve clones of each condition, either on the promoter CGIs (5’LTR + NCR CGIs considered together) or on the ETR CGI. Statistical significance was determined by unpaired T test.

Here, we studied the molecular basis to 5-AzadC reactivation potency of HIV-1 latency reversal in terms of proviral DNA demethylation. By highlighting the presence of specific epigenetic signatures in the HIV-1 promoter following 5-AzadC reactivation, we uncovered the role of the epigenetic integrator UHRF1 (Ubiquitin-like with PHD and RING finger domain 1) in the control of HIV-1 latency. As a proof-of-concept, we showed evidence that pharmacological downregulation of UHRF1 constitutes a novel therapeutic approach for anti-HIV-1 curative strategies.

## RESULTS

### 5-AzadC treatment provokes specific demethylation signatures in the HIV-1 5’LTR

We first analyzed how variations in 5-AzadC reactivation potency translated at the DNA methylation level in the HIV-1 promoter CGIs. To do so, we mock-treated or treated four clones of the CD4^+^ T-lymphoid J-Lat cell line model for HIV-1 latency with 5-AzadC. First, quantification of the viral progeny particles capsid protein p24^Gag^ in the treated culture supernatants by ELISA confirmed the variation in 5-AzadC reactivation potency *in vitro* (Fig. 1B/D/F/H, indicating a 10.6 fold, a 2.9 fold, a 2.4 fold and a 12.1 fold reactivation, for J-Lat 6.3, J-Lat 8.4, J-Lat 9.2 and J-Lat 15.4 cells, respectively). We next assessed the methylation status of the two promoter CGIs termed 5’LTR and NCR CGIs (from nt -455 to nt 179 and from nt 183 to nt 470, respectively, where nt+1 is at the U3/R junction in the 5’LTR, Fig. 1A) and of a control intragenic CGI located within *rev* termed ETR CGI (nt 7924 to nt 8196, Fig. 1A)(11). Because the two viral LTRs have identical sequences and because we wanted to specifically obtain the methylation profile of the 5’LTR, both 5’LTR and NCR CGIs were analyzed in a single amplicon. We confirmed that promoter CGIs were hypermethylated to similar levels in the four J-Lat clones in mock-treated conditions (Fig. 1C/E/G/I, 92.19%, 92.44%, 89.06% and 91.4% of 5mC, respectively, for J-Lat 6.3, J-Lat 8.4, J-Lat 9.2 and J-Lat 15.4 cells), consistent with previous observations (9, 10). Treatment with 5-AzadC provoked a global demethylation in the two promoter CGIs, though to various extents in each J-Lat clone allowing the following ranking: J-Lat 6.3 cells (29.17% of 5-AzadC-induced demethylation) > J-Lat 15.4 cells (24.99%) > J-Lat 8.4 cells (19.36%) > J-Lat 9.2 cells (5.21%) (Fig. 1, respective p-values of 0.0057; 0.0020; 0.0085; 0.0142; unpaired T test). This ranking was similar to the one we observed with the fold reactivation levels of HIV-1 production, indicating that 5-AzadC reactivation is dependent on specific demethylation of sites in the HIV-1 promoter but with a heterogeneous profile. As a control, treatment with 5-AzadC did not alter the methylation profile of the HIV-1 ETR CGI in J-Lat 6.3 cells, J-Lat 8.4 cells and J-Lat 15.4 cells (Fig. 1C/E/I, respectively). Of note and in agreement with lower basal promoter CGIs methylation level, 5- AzadC reactivation fold in HIV-1 production was the lowest in the J-Lat 9.2 clone (Fig. 1F), in which the ETR CGI was also demethylated following 5-AzadC treatment, suggesting a non-specific action for 5-AzadC on the HIV-1 promoter in this clone (Fig. 1G).

To tease out for specific regulatory mechanisms underlying the heterogeneity of 5-AzadC reactivation potency, we next mapped the probability of demethylation following 5-AzadC treatment at individual CpG positions in promoter CGIs. This probabilistic analysis highlighted that some CpGs were more prone to 5-AzadC-induced demethylation (Fig. S1 and Methods section). The most statistically significant 5-AzadC-induced differentially-demethylated positions (termed “DDMPs”) are listed in Table 1. Despite some similarities, the position of statistically significant DDMPs varied among J-Lat clones, illustrating the heterogeneity of the 5-AzadC-induced mechanisms of HIV-1 reactivation from latency recapitulated by the clones. Some DDMPS were present in sequences giving rise to viral RNA features (PBS or the packaging sequence signal ψ), suggesting a potential link between DNA methylation deposition and RNA structuration (Table 1). Importantly, several DDMPs were located within TFBS involved in HIV-1 transcriptional regulation, in the cAMP-Responsive Element (CRE)(19), NF-κB binding sites (19), Sp1 binding sites (20, 21), and interferon-stimulated response element (21) (highlighted in Figure 1A and Table 1).

**Table 1:**
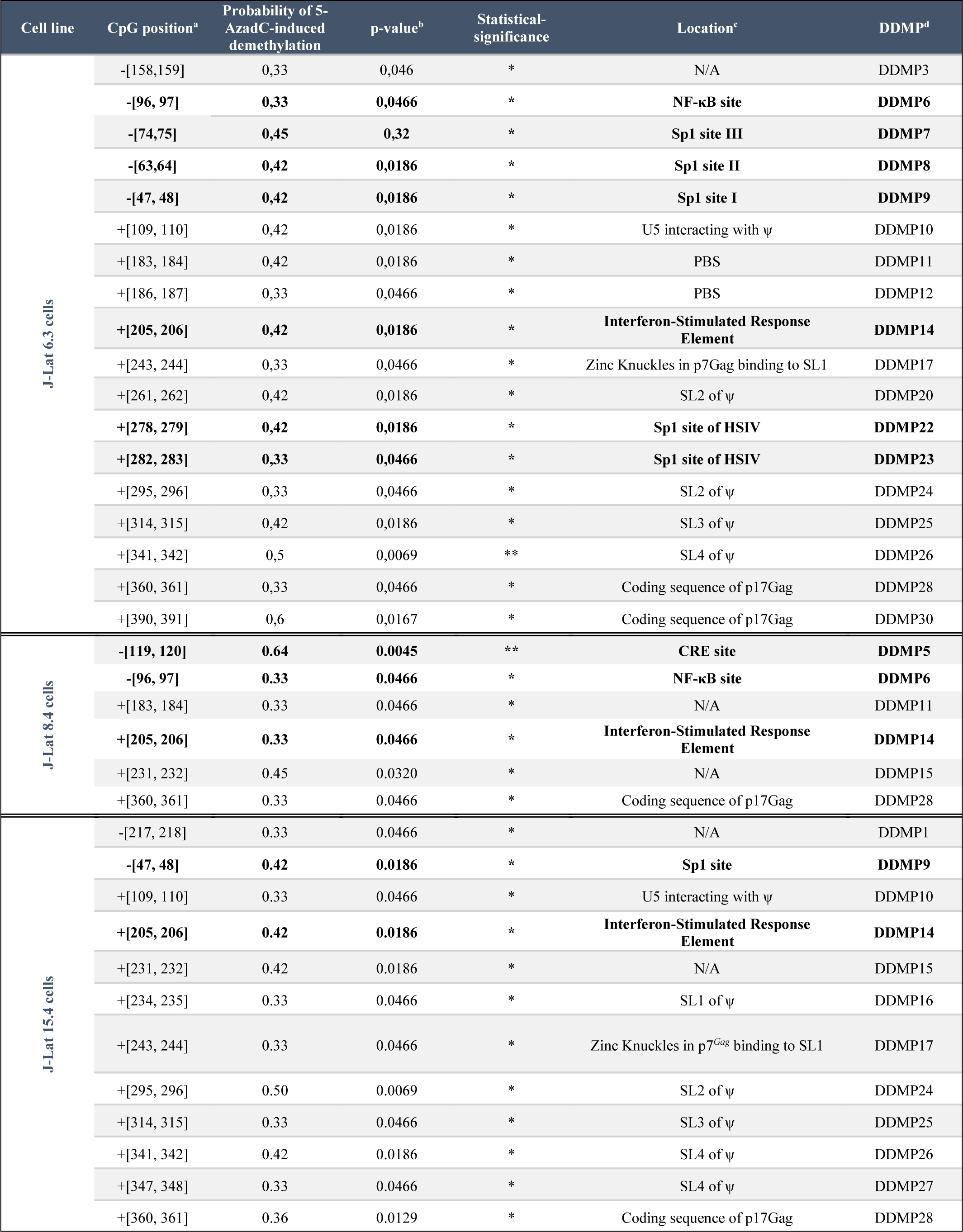
Most statistically significant 5-AzadC-induced differentially-demethylated CpG dinucleotides (DDMPs). ^a^ Position given in coordinates where nt+1 is located at the junction U3/R in the 5’LTR. ^b^ Statistical significance attributed with * for p ≤ 0.05, ** for p ≤ 0.01 and *** for p ≤ 0.001 by Fisher’s exact test. ^c^ Location of CpG within regulatory or structural elements according to the HIV-1 Database (19, 61). N/A refers to no known features. ^d^ Differentially-Demethylated Position. DDMPs located in transcription factor binding sites are in bold.

| Cell line | CpG position<sup>a</sup> | Probability of 5-AzadC-induced demethylation | p-value<sup>b</sup> | Statistical-significance | Location<sup>c</sup> | DDMP<sup>d</sup> |
| --- | --- | --- | --- | --- | --- | --- |
| J-Lat 6.3 cells | -[158,159] | 0,33 | 0,046 | * | N/A | DDMP3 |
|  | **-[96, 97]** | **0,33** | **0,0466** | **\*** | **NF-κB site** | **DDMP6** |
|  | **-[74,75]** | **0,45** | **0,32** | **\*** | **Sp1 site III** | **DDMP7** |
|  | **-[63,64]** | **0,42** | **0,0186** | **\*** | **Sp1 site II** | **DDMP8** |
|  | **-[47, 48]** | **0,42** | **0,0186** | **\*** | **Sp1 site I** | **DDMP9** |
|  | + [109, 110] | 0,42 | 0,0186 | * | U5 interacting with ψ | DDMP10 |
|  | + [183, 184] | 0,42 | 0,0186 | * | PBS | DDMP11 |
|  | + [186, 187] | 0,33 | 0,0466 | * | PBS | DDMP12 |
|  | **+ [205, 206]** | **0,42** | **0,0186** | **\*** | **Interferon-Stimulated Response Element** | **DDMP14** |
|  | + [243, 244] | 0,33 | 0,0466 | * | Zinc Knuckles in p7Gag binding to SL1 | DDMP17 |
|  | + [261, 262] | 0,42 | 0,0186 | * | SL2 of ψ | DDMP20 |
|  | **+ [278, 279]** | **0,42** | **0,0186** | **\*** | **Sp1 site of HSIV** | **DDMP22** |
|  | **+ [282, 283]** | **0,33** | **0,0466** | **\*** | **Sp1 site of HSIV** | **DDMP23** |
|  | + [295, 296] | 0,33 | 0,0466 | * | SL2 of ψ | DDMP24 |
|  | + [314, 315] | 0,42 | 0,0186 | * | SL3 of ψ | DDMP25 |
|  | + [341, 342] | 0,5 | 0,0069 | ** | SL4 of ψ | DDMP26 |
|  | + [360, 361] | 0,33 | 0,0466 | * | Coding sequence of p17Gag | DDMP28 |
|  | + [390, 391] | 0,6 | 0,0167 | * | Coding sequence of p17Gag | DDMP30 |
| J-Lat 8.4 cells | **-[119, 120]** | **0,64** | **0,0045** | **\*\*** | **CRE site** | **DDMP5** |
|  | **-[96, 97]** | **0,33** | **0,0466** | **\*** | **NF-κB site** | **DDMP6** |
|  | + [183, 184] | 0,33 | 0,0466 | * | N/A | DDMP11 |
|  | **+ [205, 206]** | **0,33** | **0,0466** | **\*** | **Interferon-Stimulated Response Element** | **DDMP14** |
|  | + [231, 232] | 0,45 | 0,0320 | * | N/A | DDMP15 |
|  | + [360, 361] | 0,33 | 0,0466 | * | Coding sequence of p17Gag | DDMP28 |
| J-Lat 15.4 cells | -[217, 218] | 0,33 | 0,0466 | * | N/A | DDMP1 |
|  | **-[47, 48]** | **0,42** | **0,0186** | **\*** | **Sp1 site** | **DDMP9** |
|  | + [109, 110] | 0,33 | 0,0466 | * | U5 interacting with ψ | DDMP10 |
|  | **+ [205, 206]** | **0,42** | **0,0186** | **\*** | **Interferon-Stimulated Response Element** | **DDMP14** |
|  | + [231, 232] | 0,42 | 0,0186 | * | N/A | DDMP15 |
|  | + [234, 235] | 0,33 | 0,0466 | * | SL1 of ψ | DDMP16 |
|  | + [243, 244] | 0,33 | 0,0466 | * | Zinc Knuckles in p7<sup>Gag</sup> binding to SL1 | DDMP17 |
|  | + [295, 296] | 0,50 | 0,0069 | * | SL2 of ψ | DDMP24 |
|  | + [314, 315] | 0,33 | 0,0466 | * | SL3 of ψ | DDMP25 |
|  | + [341, 342] | 0,42 | 0,0186 | * | SL4 of ψ | DDMP26 |
|  | + [347, 348] | 0,33 | 0,0466 | * | SL4 of ψ | DDMP27 |
|  | + [360, 361] | 0,36 | 0,0129 | * | Coding sequence of p17Gag | DDMP28 |

### Methylation of HIV-1 DDMP5 allows the recruitment of UHRF1

We next focused on the DDMP identified in J-Lat 8.4 cells and located at nucleotide positions -120 to -119 in the HIV-1 promoter, referred to as DDMP5 hereafter (Table 1, Figure 1). Indeed, DDMP5 presented the highest demethylation probability and was the most statistically significant among all identified DDMPs within TFBS in all clones (Table 1, 5-AzadC-induced demethylation probability=0.64 and p-value=0.005, Fisher’s exact test). DDMP5 is located within a known HIV-1 promoter CRE (19). Since genome-wide studies have shown that DNA methylation generally affects negatively the binding of CRE factors to their cognate sites (22), we hypothesized that DDMP5 methylation would prevent the binding of cognate transcriptional activators to the HIV-1 promoter CRE. We thus performed electrophoretic mobility shift assays (EMSAs) using radiolabeled double-stranded oligonucleotide probes containing the HIV-1NL4.3 DDMP5 sequence in an unmethylated (“DDMP5”) or in a methylated (“DDMP5-me”) form. These assays showed that the unmethylated DDMP5 probe was bound by a single retarded DNA-protein complex, termed C1 (Fig. 2A). In supershift experiments, addition of antibodies raised against CREB and CREM provoked a decreased in the complex C1 formation (Fig. 2A, lane 4 and lane 5, indicated by an asterisk), whereas addition of the IgG control or of antibodies raised against ATF1 did not affect complex formation (Fig. 2A, lane 3 and lane 6, respectively), demonstrating that the C1 complex contains both CREB and CREM proteins. Furthermore, the C1 complex was still observed when the DDMP5 probe was methylated (Fig. 2A, lane 8) and supershift experiments showed that CREB and CREM factors could bind to the same extent to the methylated and unmethylated probes (Fig. 2A, lane 10 and lane 11, respectively). We further confirmed that the binding of proteins in the C1 complex was independent of the DDMP5 methylation status, since molar excesses of both methylated and unmethylated DDMP5 competed out complex C1 formation (Fig. S2A, compare lanes 3-5 with lanes 11-13 and Fig. S2B). Together, these data indicate that DNA methylation in the HIV-1 promoter CRE site neither prevented nor decreased the binding of its cognate factors. The discrepancy between our results and the reported genome-wide DNA methylation-induced inhibition of CRE factors binding (22) could be explained by sequence differences in this TFBS motif (Fig. 2A).

**Figure 2:**
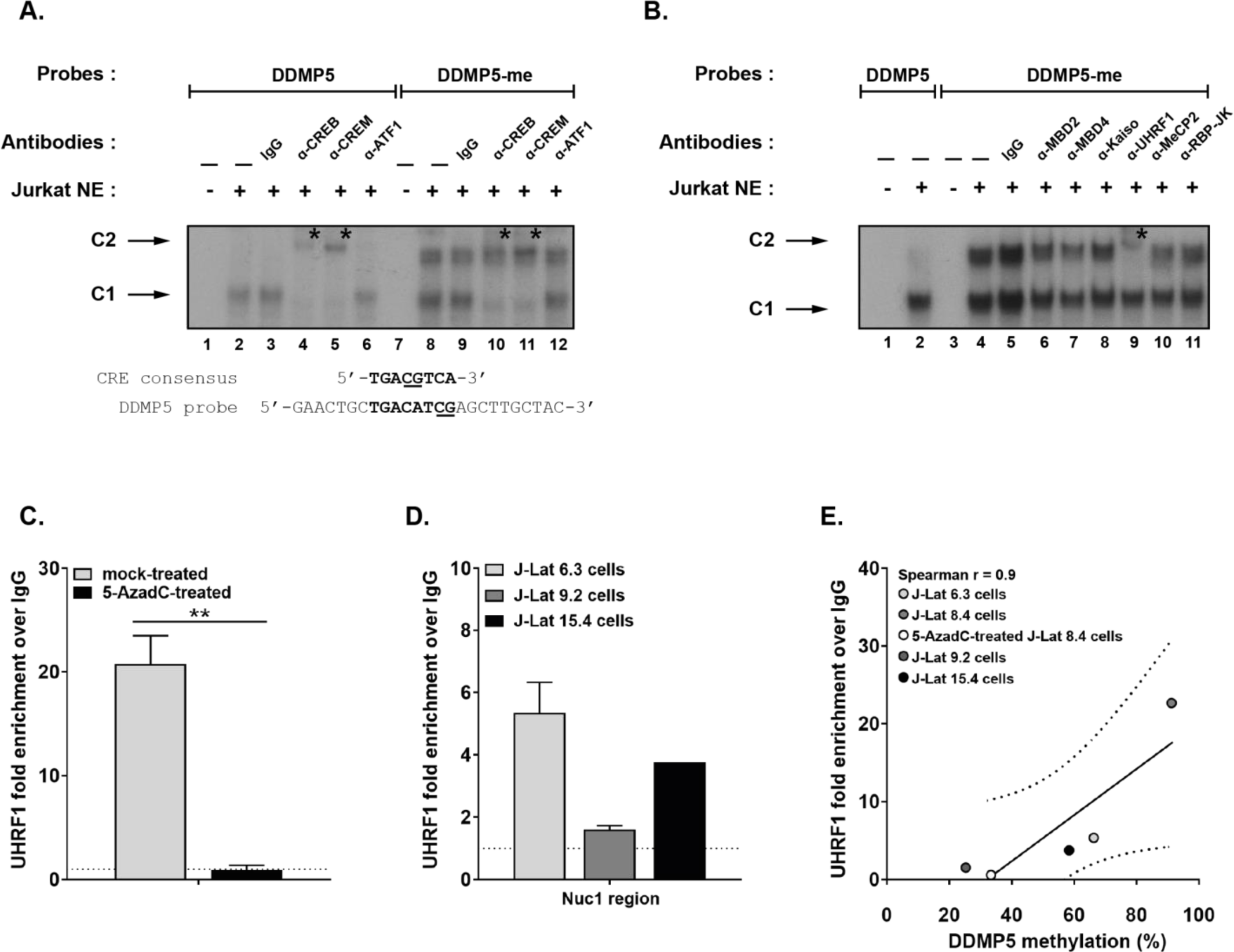
UHRF1 binds *in vitro* to the methylated DDMP5 and *in vivo* to the latent HIV-1 promoter. **(A)** The radiolabeled unmethylated or the methylated HIV-1 DDMP5 probe (respectively indicated as “DDMP5” and “DDMP5-me”) were incubated with 10µg of nuclear extracts from Jurkat T cells (“Jurkat NE”) and either with a purified rabbit IgG as a negative control (lane 3 and lane 9), or with an antibody directed against CREB/CREM family members including CREB (lane 4 and lane 10), CREM (lane 5 and lane 11) or ATF1 (lane 6 and lane 12). The figure shows the specific retarded bands of interest indicated by arrows. Supershifted complexes are indicated by asterisks. One representative experiment out of three is presented. **(B)** The “DDMP5” or “DDMP5-me” probes were incubated with 10µg of Jurkat cells NE, and either with purified rabbit IgG as a negative control (lane 5), or with an antibody directed against methylcytosines-recognizing proteins, including MBD2 (lane 6), MBD4 (lane 7), Kaiso (lane 8), UHRF1 (lane 9), MeCP2 (lane 10) and RBP-JK (lane 11). The figure shows the specific retarded bands of interest indicated by arrows. Supershifted complexes are indicated by asterisks. One representative experiment out of three is presented. **(C)** Chromatin preparations of J-Lat 8.4 cells, either mock-treated or treated with 400nM of 5-AzadC for 72h, were immunoprecipitated with an anti-UHRF1 antibody or with purified rabbit IgG, serving as a negative control. qPCRs were performed with primers hybridizing specifically to the 5’LTR, in the Nuc-1 region. Fold enrichments relative to IgG are presented, where fold enrichments for each immunoprecipitated DNA were calculated by relative standard curve on input DNA. Values represent the means of duplicate samples ± SD. Statistical significance was calculated with an unpaired T test. One representative experiment out of three is presented. **(D)** Chromatin preparation of J-Lat 6.3 cells, J-Lat 9.2 cells and J-Lat 15.4 cells were immunoprecipitated with anti-UHRF1 or purified rabbit IgG as a negative control. qPCR were performed with primers hybridizing specifically to the 5’LTR, in the Nuc-1 region. Fold enrichments relative to IgG are presented, where fold enrichments for each immunoprecipitated DNA was calculated by relative standard curve on input DNA. Values represent the means ± S.D of one representative experiment out of three. **(E)** Spearman correlation was calculated between the percentage of methylation on DDMP5 and UHRF1 recruitment fold specific to the HIV-1 promoter, based on dataset from Figure 1, Figure 2C and Figure 2D.

Interestingly, we observed by EMSAs the formation of an additional retarded complex, termed C2, with the methylated HIV-1 DDMP5 probe (Fig. 2A, lanes 8 to 12). We found that the C2 complex was formed only when the DDMP5 probe was methylated and was competed out only by methylated DDMP5 oligonucleotides (Fig. S2A, lanes 7-9). Furthermore, formation of the C2 complex was not competed out by molar excesses of the methylated consensus for methyl-binding domain (MBD) proteins, or of the methylated consensus for Sp1, indicating that the proteins contained within the C2 complex were specific to the methylated HIV-1 DDMP5, and not to any 5mC-containing sequence (Fig. S2C). Because our *in vitro* experiments showed that, rather than preventing the binding of transcriptional activators, DDMP5 methylation allowed the binding of methylCpG-recognizing proteins to the HIV-1 promoter, we investigated the nature of the proteins present in the C2 complex. To do so, we performed additional supershift experiments, using antibodies raised against proteins known to bind methylcytosines (MBD2, MBD4, MeCP2, Kaiso, UHRF1 and RBP-JK)(23, 24). Addition of an antibody raised against UHRF1, but not against the other proteins, altered the formation of the C2 complex, concomitantly with the appearance of a supershifted complex of lower mobility (Fig. 2B, lane 9, indicated with an asterisk), while addition of IgG did not affect complex C2 formation (Fig. 2B, lane 5). These results thus indicated that the C2 complex contained UHRF1 that bound *in vitro* to the methylated HIV-1 DDMP5.

To demonstrate *in vivo*, within the context of chromatin, the relevance of UHRF1 binding to the HIV-1 promoter, we performed chromatin immunoprecipitation (ChIP) assays in J-Lat 8.4 cells. Using primers hybridizing specifically to the HIV-1 5’LTR, in the Nuc-1 region, we showed that UHRF1 was recruited to the latent promoter in J-Lat 8.4 cells (Fig. 2C, 20.75-fold recruitment). Furthermore, 5- AzadC-induced demethylation, allowing DNA demethylation of DDMP5 (Fig. 1E), was accompanied by a statistically significant decrease in UHRF1 recruitment to the viral promoter (Fig. 2C, 22.9-fold decrease, p=0.01, unpaired T test). As a control, we quantified *UHRF1* mRNA and protein levels in J-Lat 8.4 cells in response to 5-AzadC (Fig. S3A and Fig. S3B, respectively), and showed that 5-AzadC did not alter UHRF1 expression, thus demonstrating a direct link between UHRF1 decreased recruitment to the HIV-1 promoter and its 5-AzadC-induced demethylation. Despite the fact that DDMP5 had been identified in the J-Lat 8.4 clone, we next assessed whether UHRF1 *in vivo* recruitment to the HIV-1 5’LTR was a common feature of HIV-1 latency. These additional ChIP experiments showed that UHRF1 was also recruited to the latent viral promoter in the J-Lat 6.3 and J-Lat 15.4 cells, albeit to lower levels than in the J-Lat 8.4 cells (Fig. 2D, 5.35-fold and 3.75-fold recruitment, respectively). To determine how UHRF1 recruitment modalities correlated to DDMP5 methylation status in the 5’LTR, we plotted UHRF1 fold recruitment from our ChIP results to DDMP5 methylation level in latent conditions in all four J-Lat clones. This showed that UHRF1 *in vivo* recruitment followed the status of the DDMP5 methylation status in the four clones analyzed, with a trend towards higher recruitment of UHRF1 to more methylated DDMP5 (Fig. 2E). For instance, J-Lat 9.2 cells, in which DDMP5 was already largely unmethylated in basal conditions (Fig. 1G, 9 clones unmethylated on 12) showed weak *in vivo* UHRF1 recruitment to the HIV-1 promoter (Fig. 2E). Of note, the lack of significant correlation between UHRF1 *in vivo* recruitment and the DDMP5 methylation status in the 5’LTR can be explained by its independent recruitment to the viral promoter through other epigenetic marks, thanks to UHRF1 domains for histone methylation or acetylation (25, 26), through interaction with other epigenetic enzymes (27, 28) or through direct recognition of binding motifs in the 5’LTR (29).

Taken together, our results demonstrated that the 5’LTR DDMP5 position corresponds to an *in vitro* binding site for UHRF1. We confirmed the *in vivo* recruitment of UHRF1 to the latent HIV-1 5’LTR and showed it is proportional to the methylation level of a single CpG residue. Furthermore, reactivation of HIV-1 gene expression following 5-AzadC treatment was accompanied by the demethylation of DDMP5 and by a decreased UHRF1 recruitment to the viral promoter, suggesting a role for UHRF1 in DNA methylation-mediated silencing of HIV-1 gene expression.

### UHRF1 transcriptional repression of the HIV-1 promoter is DNA methylation-dependent

To determine the involvement of UHRF1 in the maintenance or establishment of transcriptional silencing at the HIV-1 promoter during latency, we induced the downregulation of endogenous UHRF1 by shRNAs. J-Lat 8.4 cells were mock-transduced, or stably transduced with lentiviral vectors expressing the puromycin-resistance gene and containing one out of four different shRNAs targeting UHRF1 mRNA (shUHRF1#1-4) or a control non-targeting shRNA (shNT). UHRF1 knockdown in selected puromycin-resistant clones was confirmed both by western blot analyses and by RT-qPCR (Fig. S4A and S4B, respectively). Because UHRF1 is essential in the cell cycle control and its downregulation is associated with cellular mortality (30, 31), we selected for further analyses one shUHRF1 that did not provoke the most efficient downregulation of UHRF1 but that would, therefore, not be counter-selected (Fig. S4, pLV shUHRF1#4). First, by ChIP experiments, we showed that RNAPII recruitment to the Nuc-1 region of the HIV-1 promoter was statistically higher in UHRF1-depleted J-Lat 8.4 cells than in shNT-transduced cells, consistent with a release of viral transcriptional blocks from latency (Fig. 3A, 2.48-fold increase, p=0.008, unpaired T test). Reactivation of HIV-1 gene expression from latency in J-Lat 8.4 cells following UHRF1 knockdown was further studied by quantifying by RT-qPCR initiated (TAR region) and elongated (*tat* region) HIV-1 transcripts (Fig. 3B). A statistically higher number of initiated and elongated transcripts was observed when UHRF1 was knocked down in J-Lat 8.4 cells, compared to the amount measured in shNT-transduced cells (Fig. 3B, 3.29-fold increase, p=0.0011 and 1.7-fold, p=0.034, for TAR and *tat*, respectively, unpaired T test). This indicates that the observed increase in RNAPII recruitment was accompanied by an increased transcription initiation and elongation from the HIV-1 promoter. In addition, we observed a statistically significant increase in multiply spliced (MS) HIV-1 RNA (32) in shUHRF1-transduced cells compared to shNT-transduced cells (Fig. 3B, 2.99- fold increase, p=0.0004, unpaired T test). Finally, quantification of p24^Gag^ capsid protein by ELISA in culture supernatants from puromycin-resistant clones showed that UHRF1 knockdown was accompanied by a higher HIV-1 production than the one observed in the shNT-transduced cells (Fig. 3C, 4.5-fold increase, p=0.02, unpaired T test). We observed this statistically significant increase in HIV-1 production upon UHRF1 depletion with the four shUHRF1 we used (Fig. S4C), strengthening the specific role of UHRF1 in HIV-1 silencing. Of note, J-Lat 8.4 cells transduction with the control, non-targeting shRNA, also caused reactivation of HIV-1 gene expression and production, though to lower levels than transduction of the shUHRF1, as seen by increased HIV-1 initiating, *gag* and multiply spliced transcripts (Fig. 3B) and increased HIV-1 production (Fig. 3C and Fig. S4C). This was consistent with the use of lentiviral shRNA vectors and was in agreement with a previous report (33). Nevertheless, we observed the reactivation of HIV-1 gene expression and production when shUHRF1-transduced conditions were normalized to shNT-transduced conditions, and *a fortiori*, when normalized to mock-transduced conditions (Fig. 3A-C). Together, these data demonstrated that UHRF1 knockdown enables a full release (i.e. up to the completion of the replication cycle and the production of progeny particles) of the transcriptional blocks from latency in J-Lat 8.4 cells, thereby indicating a role for UHRF1 in the maintenance of HIV-1 latency.

**Figure 3:**
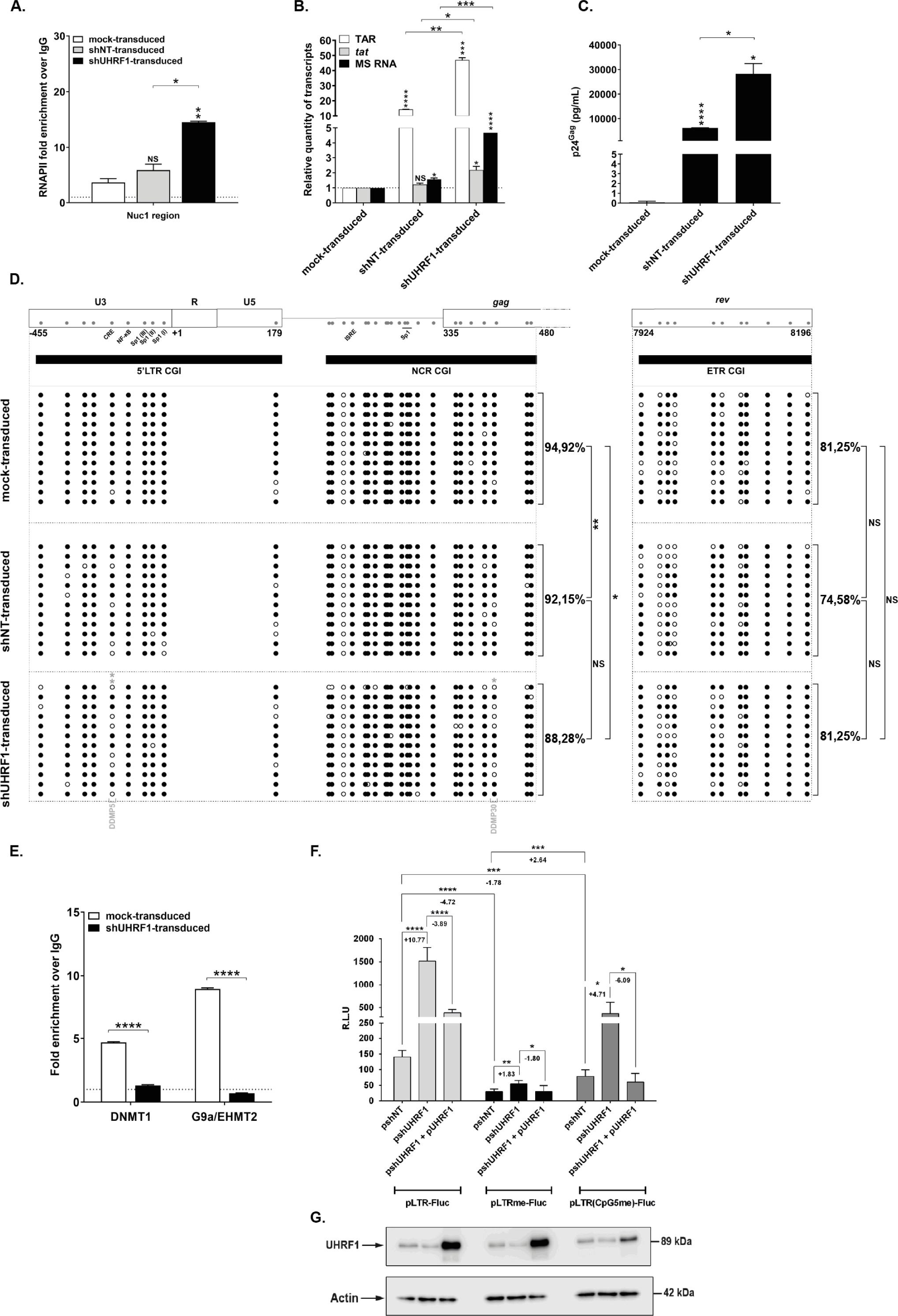
UHRF1 silencing of HIV-1 transcription is dependent on 5’LTR methylation. **(A)** Chromatin was prepared from J-Lat 8.4 cells, either mock-transduced, control-transduced (with non- targeting shRNA, indicated as “shNT-transduced”) or shUHRF1-transduced (indicated as “shUHRF1- transduced”). Immunoprecipitations were performed using anti-RNAPII or purified rabbit IgG, as a negative control. qPCRs were performed with primers hybridizing specifically to the 5’LTR, in the Nuc-1 region. Fold enrichments relative to IgG are presented, where fold enrichments for each immunoprecipitated DNA were calculated by relative standard curve on input DNA. Values represent the means of duplicate samples ± SD. **(B)** Total RNA preparations from mock-transduced, shNT- transduced or shUHRF1-transduced J-Lat 8.4 cells were reverse transcribed. Initiated (TAR region), elongated (*tat* region) transcripts, or HIV-1 multiple-spliced RNA (MS RNA) were quantified by RT- qPCR using GAPDH as first normalizer and the mock-transduced condition as second normalizer. Means from duplicate ± SD are indicated. **(C)** Cultures supernatants from mock-transduced, shNT- transduced or shUHRF1-transduced J-Lat 8.4 cells were probed for viral production as measured by ELISA on p24^Gag^ capsid protein. **(D)** DNA methylation mapping performed by sodium bisulfite sequencing is presented for the promoter CGIs or the intragenic ETR CGI in J-Lat 8.4 cells mock- transduced, shNT-transduced or shUHRF1-transduced, as indicated. Unmethylated and methylated CpG dinucleotides are respectively represented with open and closed circles, where each line corresponds to individual sequenced molecules. The global methylation level presented correspond to mean percentages of methylated CpGs for the twelve clones of each condition, either for the promoter CGIs considered together (5’LTR + NCR CGIs) or for the ETR CGI. Panel A to D originate from the same representative experiment out of three. **(E)** Chromatin was prepared from J-Lat 8.4 cells, either mock- transduced or shUHRF1-transduced. Immunoprecipitations were performed using anti-DNMT1, anti- G9a or purified rabbit IgG, as a negative control. qPCRs were performed with primers hybridizing specifically to the 5’LTR, in the Nuc-1 region. Fold enrichments relative to IgG are presented, where fold enrichments for each immunoprecipitated DNA were calculated by relative standard curve on input DNA. Values represent the means of duplicate samples ± SD. **(F)** HEK293T cells were transfected either with 600ng of the pshNT vector or with 600ng of the pshUHRF1#4. Twenty-four hours after this initial transfection, 400ng of the pLTR-Fluc, the pLTRme-Fluc or the pLTR’Cpg5me)-Fluc reporter constructs together with or without 200ng of the plasmid overexpressing UHRF1 (pUHRF1) were co-transfected. Luciferase activities were measured in the cell lysates after further 24h post-transfection. Results are presented as histograms of “relative luciferase units” (R.L.U.), corresponding to the Fluc activiy normalized to the total levels of proteins. Means and standard errors of triplicate samples are represented. An experiment representative of three independent experiments is shown. Statistical significance was assessed by an unpaired T test. **(G)** UHRF1 and β-actin, serving as a loading control, protein levels were assessed by immunoblot in cell lysates of the corresponding transfection points.

**Figure 4:**
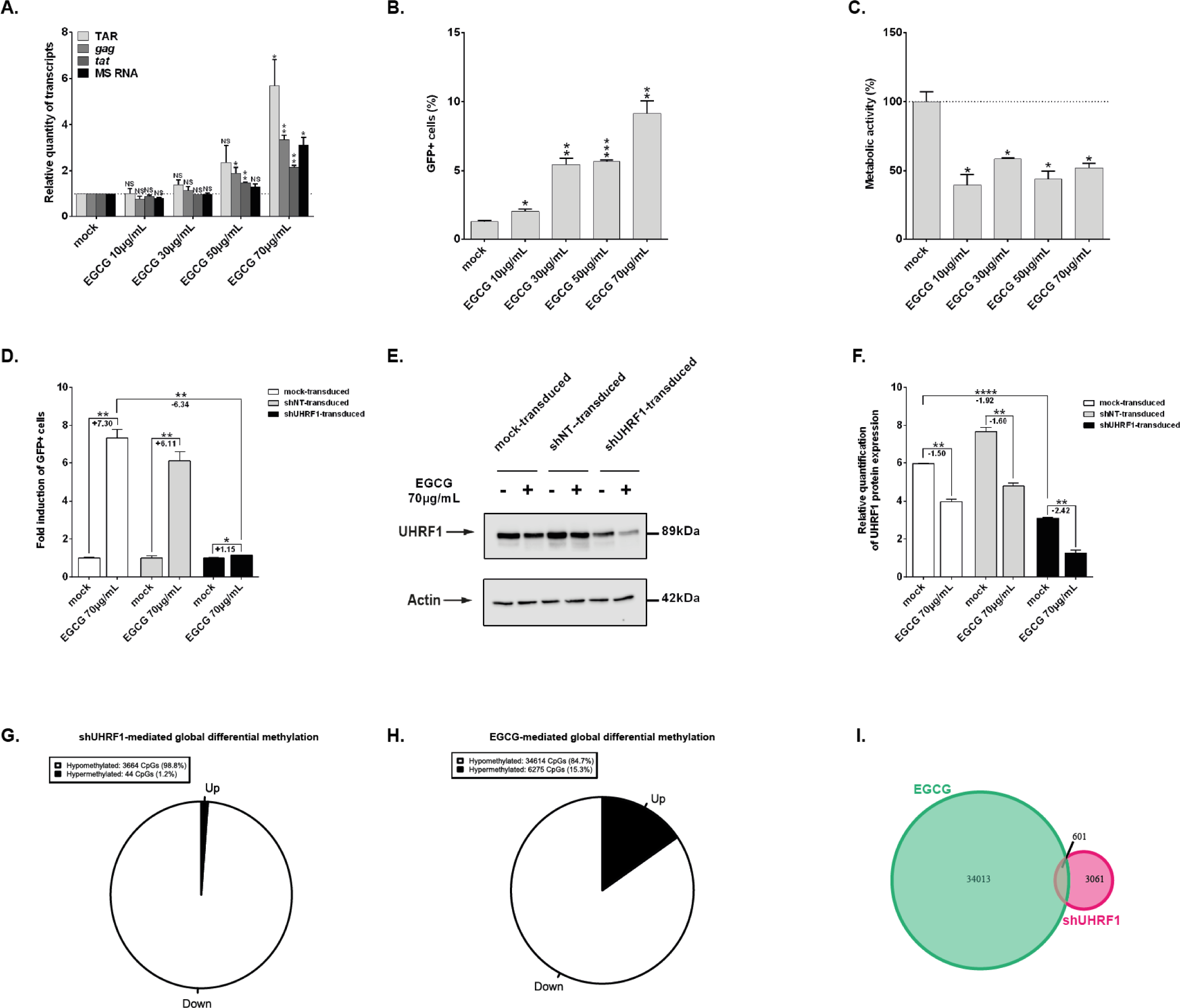
HIV-1 transcription is reactivated from latency upon UHRF1 downregulation by epigallo-catechin-3-gallate. **(A)** Total RNA preparations from J-Lat 8.4 cells mock-treated or treated with increasing doses of EGCG for 24h were used in RT-qPCR to quantify initiated (TAR region), *gag*, elongated (*tat* region) transcripts, or HIV-1 MS RNA, using GAPDH as normalizer. **(B)** After 24h of EGCG increasing doses treatment, J-Lat 8.4 cells were analyzed by flow cytometry to quantify the percentage of GFP+ cells. **(C)** WST-1 proliferation assay, reflective of metabolic activity, was performed on J-Lat 8.4 cells treated for 24h with increasing doses of EGCG. The result obtained with mock-treated cells was set a value of 100%. A, B and C originate from the same representative experiment out of three. Means ± SD of duplicates are presented. Statistical significance was calculated with an unpaired T test and corresponds to comparisons to the mock-treated condition. **(D)** J-Lat 8.4 cells, either mock-transduced or shUHRF1- transduced were treated for 24h with 70µg/mL of EGCG. The percentage of GFP positive cells was assessed by flow cytometry and was normalized to the respective mock conditions. Means ± SD of duplicates representative of three independent experiments are presented. Statistical significance was assessed by an unpaired T test. **(E)** Whole protein levels of the experiments presented in Fig. 4D were loaded and probed for the presence of UHRF1 and and β-actin, serving as a loading control. **(F)** Quantification of the western blot presented in Fig. 4E was performed in ImageJ. Means ± SD of two independent quantifications are presented. Statistical significance was assessed by an unpaired T test. **(G)** An Infinium assay with gDNA from above was conducted. Differential methylated CpGs upon EGCG treatment were plotted on a pie chart, where the black slice represents hypermethylated CpGs and the white slice represents hypomethylated CpGs. **(H))** An Infinium assay with gDNA from Fig. 3 was conducted. Differential methylated CpGs upon shUHRF1 transduction were plotted on a pie chart, where the black slice represents hypermethylated CpGs and the white slice represents hypomethylated CpGs. **(I)** Venn diagram representing the number of hypomethylated CpGs in both conditions is presented.

Because UHRF1 is an important epigenetic integrator in the heterochromatinization of *cis*- regulatory sequences (25, 34, 35), we next further investigated how UHRF1 knockdown and subsequent reactivation of HIV-1 gene expression translated in terms of DNA methylation modifications on the HIV-1 promoter. UHRF1 knockdown was associated with a statistically significant decrease in global HIV-1 promoter DNA methylation compared to mock-transduced conditions (Fig. 3D), whereas no DNA demethylation was observed on the control ETR CGI. These results thus indicate that UHRF1 depletion leads to HIV-1 transcriptional reactivation through specific 5’LTR demethylation. Of note, the demethylation signatures in the HIV-1 promoter CGIs in response to UHRF1 depletion were similar than those in 5-AzadC-treated cells (compare Fig. 3D and Fig. 1E). Indeed, the fifth CpG dinucleotide in the 5’LTR CGI showed a high demethylation probability in response to UHRF1 knockdown (Fig. S4), linking mechanistically UHRF1 with 5-AzadC-induced reactivation in J-Lat 8.4 cells. In addition, shNT transduction that provoked HIV-1 reactivation was also accompanied by global 5’LTR demethylation and by demethylation of DDMP5, further highlighting that the DDMP5 methylation state is functionally linked to the level of HIV-1 gene silencing (Fig. 3D). UHRF1 has been showed to interact with multiple epigenetic enzymes, including DNMT1 (30, 36) and G9a/EHMT2 (27, 28), that are important actors in the HIV-1 promoter heterochromatinization during latency (reviewed in (8)). We thus determined the effect of UHRF1 downregulation on the *in vivo* recruitment of DNMT1 and G9a/EHMT2 to the viral promoter. Following depletion of UHRF1 and reactivation of HIV-1 gene expression and production from latency, a significant decrease in DNMT1 and in G9a/EHMT2 was observed on the viral promoter (Fig. 3E, 3.61-fold decrease, p=0.0003 and 12.71-fold decrease, p=0.00005, unpaired T test, for DNMT1 and G9a, respectively). These results thus indicate that UHRF1 silences HIV-1 gene expression during latency by actively promoting the accumulation of DNA methylation on the viral promoter *via* its recruitment of DNMT1. Furthermore, we demonstrate that UHRF1 also recruits G9a/EHMT2 to the latent promoter, thereby linking DNA methylation with the accumulation of repressive histone methylation.

UHRF1 possesses multiple repression mechanisms of gene expression and our data suggest that it can be recruited to the 5’LTR independently of DDMP5 methylation. We thus further dissected the dependency of UHRF1 to DNA methylation and in particular, to the methylation status of DDMP5 for its role in HIV-1 transcriptional repression. To do so, we subcloned the HIV-1 5’LTR region in a reporter construct, where the LTR controls the firefly luciferase gene and is either unmethylated (referred as to the “pLTR-FLuc” vector), in a hypermethylated state (i.e. where each CpG of the 5’LTR has been artificially methylated, the resulting construct being termed “pLTRme-Fluc”), or where only the 5^th^ CpG dinucleotide corresponding to the DDMP5 of the LTR is methylated (referred as to the “pLTR(CpG5me)-Fluc” vector). First, these three reporter vectors were transiently transfected in HEK293T cells along with the control non-targeting shRNA vector (referred as to the “pshNT”). These transfections showed that the pLTRme-Fluc vector presented a statistically significant decreased luciferase activity in comparison to the pLTR-Fluc vector (Fig. 3F, 4.72-fold decrease, p<0.0001, unpaired T test), confirming that methylation of the LTR provoked a decrease of its promoter activity. Interestingly, methylation of the DDMP5 position alone was sufficient to reduce the LTR promoter activity in a statistically relevant manner (Fig. 3F, 1.78-fold decrease, p=0.0003, unpaired T test), albeit not as much as with the fully-methylated LTR (Fig. 3F, 2.64-fold increase, p=0.0003, unpaired T test). These data confirmed the importance of DDMP5 methylation in recapitulating DNA methylation-mediated repression on the HIV-1 promoter. Second, we transiently co-transfected the reporter LTR constructs along with the UHRF1-targeting shRNA vector (“pshUHRF1”). Downregulation of endogenous UHRF1 in HEK293T cells was confirmed by western blot (Fig. 3G) and was associated with statistically significant increased luciferase activities for all three constructs (Fig. 3F, 10.77-fold increase, p<0.0001 ; 1.83-fold increase, p=0.0014 and 4.71-fold increase, p=0.0152, for the pLTR-Fluc, pLTRme-Fluc and pLTR(CpG5me)-Fluc constructs, respectively and according to an unpaired T test), confirming the repressive role of UHRF1 in the control of HIV-1 gene expression. Interestingly, in this reporter system, UHRF1-mediated repression of the HIV-1 promoter activity was proportionally lower when the LTR was totally methylated or methylated on the DDMP5 in comparison to the repression observed when the LTR was unmethylated (Fig. 3F, compare the 1.83-fold and 4.71-fold increases with the 10.77-fold increase, for the fully-methylated, DDMP5-methylated and unmethylated LTRs, respectively). These results indicated that UHRF1 repressive activity was dependent on the HIV-1 promoter DNA methylation status, and in particular, that this repression was partially, but not totally, dependent on the DDMP5 methylation status. To confirm the dependency of UHRF1 on DNA methylation for its role in HIV-1 transcriptional repression, we performed a rescue experiment in which we transiently co-transfected reporter constructs along with the UHRF1-targeting shRNA vector then, after twenty-four hours, we added an UHRF1 expression vector (“pUHRF1”) before assaying the luciferase activities after another twenty-four hours. Overexpression of UHRF1 was confirmed by western blot (Fig. 3G) and decreased the luciferase activities of all three reporter vectors in comparison to the conditions of UHRF1 downregulation (Fig. 3F, 3.88-fold decrease, p<0.0001 ; 1.80-fold decrease p=0.0217 and 6.09-fold decrease, p=0.0114 for the pLTR-Fluc, pLTRme-Fluc and pLTR(CpG5me)- Fluc constructs, respectively), thereby confirming the specific role of UHRF1 in the repression of HIV- 1 promoter activity. In particular, this effect was proportionally more important for the DDMP5- methylated construct, then for the fully-methylated construct, in comparison to the unmethylated LTR construct (Fig. 3F, compare the 6.09-fold and 1.80 -fold to the 3.89-fold, for the DDMP5-methylated, fully-methylated and unmethylated LTRs, respectively). Together, these results indicated that in the context of an *in vitro* HIV-1 5’LTR reporter system, UHRF1 repression of viral transcription depended in part but not exclusively on DNA methylation of the viral promoter. Importantly, our system allowed to specifically dissect the contribution of the single DDMP5 methylation status, showing its role in the repression of HIV-1 promoter activity.

Altogether, our results demonstrated the DNA methylation-mediated role of UHRF1 in the HIV- 1 promoter silencing during latency.

### Pharmacological downregulation of UHRF1 by EGCG promotes HIV-1 reactivation from latency

Understanding the molecular mechanisms of HIV-1 latency has allowed the development of several classes of LRAs (5). Our results on UHRF1 positioned this cellular factor as an attractive pharmacological target for HIV-1 latency reversal strategies. Epigallocatechin-3-gallate (EGCG), the major polyphenolic compound of green tea, has been shown to downregulate UHRF1 expression (37). Accordingly, we showed that increasing concentrations of EGCG steadily decreased UHRF1 protein levels starting from 30µg/mL of EGCG in J-Lat 8.4 cells (Fig. S5A). This protein level decrease was not accompanied by a decrease in *UHRF1* mRNA level, as quantified by RT-qPCR (Fig. S5B), consistent with a previous report showing that EGCG targets UHRF1 proteins but not *UHRF1* transcripts (37).

To assess the LRA potential of EGCG *in vitro*, we quantified HIV-1 transcripts by RT-qPCR in treated J-Lat 8.4 cells (Fig. 4A). We observed statistically significant increases in initiated (TAR region), elongated (*gag* and *tat* regions), and MS HIV-1 transcript levels in EGCG-treated compared to mock-treated conditions (Fig. 4A, 5.70-fold and p=0.03, 3.35-fold and p=0.003, 2.15-fold and p=0.003, and 3.11-fold increase and p=0.01 for TAR, *tat*, *gag* and MS RNA, respectively at 70µg/mL of EGCG). Furthermore, quantification of GFP^+^ cells by flow cytometry showed that starting from 10µg/mL of EGCG, a release of the post-transcriptional blocks to the production of HIV-1 was observed (Fig. 4B). In addition, the cellular metabolic activity in J-Lat 8.4 cells after treatment with increasing EGCG doses was decreased in a statistically relevant manner, with a metabolic activity of 36% observed at the highest EGCG dose (Fig. 4C). Despite increased levels of *gag* transcripts (Fig. 4A), EGCG did not reactivate HIV-1 protein production in J-Lat 8.4 cells, as measured by p24^Gag^ capsid protein ELISA in cell supernatants (Fig. S5C). These data were consistent with a previous report indicating that EGCG destabilizes HIV-1 particles by binding to envelope phospholipids, thereby inducing their deformation (38).

**Figure 5:**
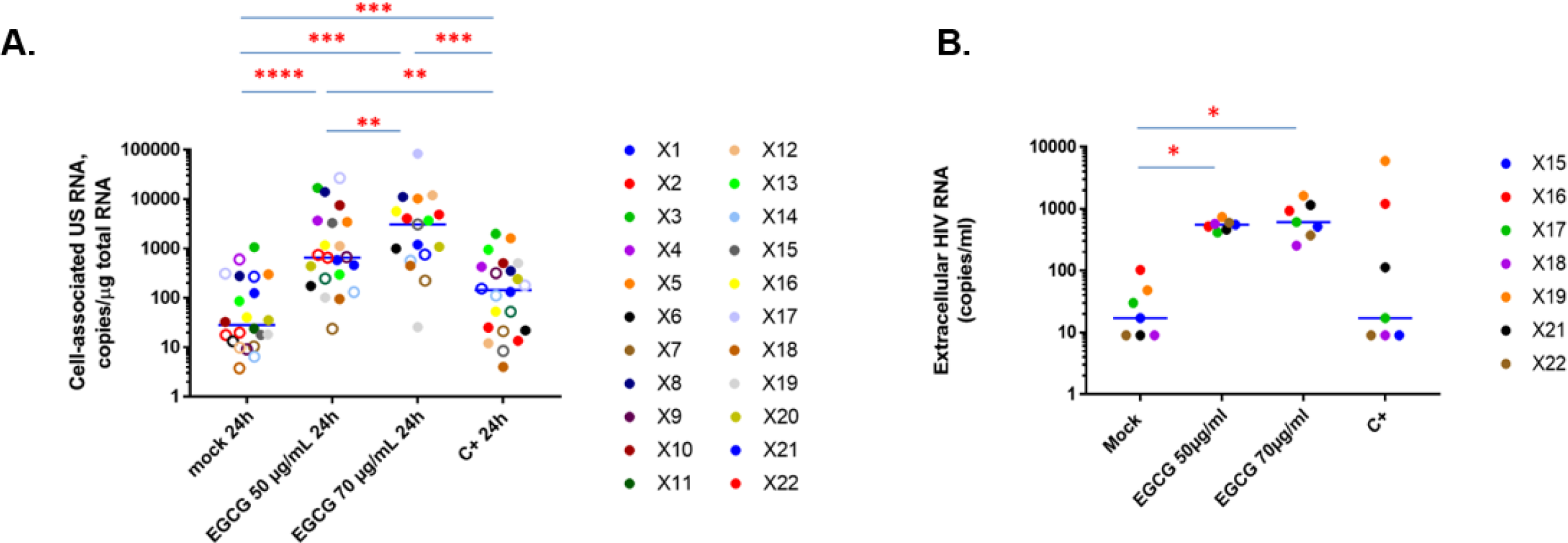
EGCG reactivates the expression of viral RNA in *ex vivo* cultures of CD8^+^-depleted PBMCs isolated from cART-treated aviremic HIV-1+ individuals. *Ex vivo* cultures of CD8^+^-depleted PBMCs isolated from 22 cART-treated aviremic HIV^+^ individuals were mock-treated or treated for 24h with EGCG, at the indicated concentrations, or with anti-CD3+anti-CD28 antibodies serving as positive control stimulation. **(A)** Total intracellular RNA was extracted and cell-associated HIV-1 US RNA was quantified. Medians are represented. Open circles depict undetectable values, censored to the assay detection limits. The latter depended on the amounts of input cellular RNA and therefore differed between samples. Statistical significance was determined by paired Wilcoxon tests, where pairs were included in the analysis only when either (i) both values in a pair were detectable, or (ii) one value in a pair was undetectable and the other detectable, and the maximal value of the undetectable (the assay detection limit) was lower than the detectable. **(B)** In 7 out of 22 HIV^+^ individuals, the concentration of HIV-1 extracellular genomic RNA in culture supernatants was also determined (in copies/ml).

Since EGCG is a broad-acting compound (39), we next investigated EGCG modes of action in HIV-1 latency reversal, specifically, their dependency on UHRF1 downregulation. First, we performed EGCG reactivation assays in latently-infected J-Lat 8.4 cells in which UHRF1 expression had been downregulated. By normalizing each EGCG treatment to its respective mock control, we observed a statistically-significant decrease in EGCG reactivation potency in shUHRF1-transduced versus mock-transduced cells (Fig. 4D, 6.34-fold decrease, p=0.0032, unpaired T test), whereas UHRF1 expression was further downregulated in EGCG-treated shUHRF1-transduced cells in comparison to mock-treated, mock-transduced cells (Fig. 4E and Fig. 4F). Second, we assessed the DNA methylation signatures occurring at genome-scale while knocking down UHRF1 or treating latently-infected cells with EGCG and compared them. To do so, we performed an Infinium Human Methylation 850K array (40). By using unsupervised analyses, such as hierarchical clustering and principal component analyses, we revealed a strong effect of both EGCG treatment and UHRF1 knockdown on the cellular methylome, with treated samples being distinctly different from control samples (Fig. S6A and Fig. S6B). In addition, EGCG treatment and UHRF1 knockdown methylation profiles partially clustered together, suggesting that the effect of the two conditions on the DNA methylome is only partly similar. We identified 3664 hypomethylated CpGs through UHRF1 knockdown and 34614 hypomethylated CpGs through EGCG treatment (Fig. 4G and Fig. 4H, respectively). As already suggested by the unsupervised analyses, the overlap between differential CpGs through EGCG treatment and UHRF1 knockdown, while small, was statistically significant (Fig. 4I, 601 sites, hypergeometrical p-value < 1e-170), suggesting that some but not all mechanisms involved in the two processes were similar. We further assessed which pathways were affected at the DNA methylation level in EGCG-treated and shUHRF1-transduced conditions using a Gene Set Enrichment Analysis (GSEA)(41). This analysis further confirmed the partial but not total overlap between the two conditions (Fig. S6C and S6D).

Together, these experiments indicated that EGCG reactivates HIV-1 from latency in part *via* the downregulation of UHRF1, although, this compound has a broader reactivation capacity on HIV-1 gene expression, in line with its pleiotropic action on the HIV-1 replication cycle (38, 42).

### EGCG induces HIV-1 expression in CD8^+^-depleted PBMCS from HIV-1^+^ aviremic individuals

The development of LRAs has been guided by deciphering HIV-1 latency molecular mechanisms in *in vitro* cell models (5). However, these models do not completely recapitulate the biological properties of *in vivo* latent reservoirs (43). Therefore, we next evaluated the LRA potency of EGCG *ex vivo*, using cultures of CD8^+^-depleted PBMCs isolated from blood of cART-treated aviremic HIV-1^+^ individuals.

We first assessed cellular viability and metabolic activity in *ex vivo* cultures of CD8^+^-depleted PBMCs from six healthy donors in response to EGCG treatments (Fig. S7). Neither TCR stimulation, serving as a positive control, nor increasing EGCG doses affected cellular viability (Fig. S7A). However, consistent with our observations in J-Lat cells, metabolic activity was affected by EGCG, although the median of metabolic activity remained tolerable (Fig. S7B). Suitable LRA candidates for anti-HIV-1 latency strategies *in vivo* should limit non-specific or strong immune T-cell activation. Therefore, we assessed the level of cell surface activation markers HLA-DR (late activation marker), CD25 (intermediate activation marker), CD69 (early activation marker) and CD38 (late activation marker and predictor of HIV-1 progression) in comparison to the mock-treated condition (Fig. S7C-F). TCR stimulation consistently and statistically increased the levels of each marker, while EGCG treatment increased slightly but statistically the surface expression of CD69 (Fig. S7E). Because this increase in CD69 expression was not associated with later cell activation when EGCG stimulation was sustained for 6 days (data not shown), we attributed it to an indirect epigenetic effect of EGCG (44). Flow cytometry analyses also highlighted that increasing doses of EGCG were associated with a statistically significant decrease in CD4 expression on the treated CD8^+^-depleted PBMCs (Fig. S7G), which would, in the context of HIV-1 infection, reduce the number of target cells and therefore limit HIV-1 dissemination.

Based on these observations of tolerable cytotoxic, metabolic and immune effects, we next investigated HIV-1 recovery in CD8^+^-depleted PBMCs in response to EGCG treatments. To do so, we purified CD8^+^-depleted PBMCs from 22 HIV^+^ aviremic cART-treated individuals (Table S1A) and evaluated the frequency of infected cells during plating by quantification of cell-associated total HIV-1 DNA (Table S1B). *Ex vivo* cultures were then mock-treated, treated with 50µg/mL or 70µg/mL of EGCG or activated with anti-CD3 + anti-CD28 antibodies as a positive control. Because of EGCG effect in degrading viral particles, reactivation of HIV-1 gene expression was measured unequivocally by the quantification of intracellular HIV-1 RNA. These quantifications showed that EGCG potently increased HIV-1 unspliced RNA levels in patient cells, even to higher levels than TCR activation (Fig. 5A). Moreover, HIV-1 US RNA/DNA ratios were statistically increased in EGCG-treated conditions (Fig. S8A and S8C), indicating that proviruses were more transcriptionally active. Quantification of HIV-1 extracellular RNA in supernatants, serving as a surrogate for the completion of the viral replication cycle, further indicated a statistically significant increase in HIV-1 extracellular RNA when CD8^+^- depleted PBMCs from HIV^+^ individuals were submitted to EGCG treatments (Fig. 5B). Thus our result show that, by destabilizing HIV-1 particles in reactivated cell cultures (38), EGCG treatment released HIV-1 RNA from virions *ex vivo*. Accordingly, we observed a statistically-significant increase in HIV- 1 extracellular RNA/DNA ratios in EGCG-treated compared to mock-treated *ex vivo* patient cell cultures (Fig. S8B and S8C), indicating that not only transcriptional but also post-transcriptional latency blocks were overcome by EGCG treatment. Altogether, our results highlight the strong potency of EGCG as a new LRA *ex vivo* allowing to reactivate HIV-1 transcription to the completion of the replication cycle, while maintaining low immune activation level, and even allowing to prevent *de novo* infections by decreasing cell surface CD4 marker expression and by degrading reactivated HIV-1 particles.

Finally, because our *in vitro* data pointed to the pleiotropic action of EGCG in its HIV-1 reactivation capacity, we further assessed the contribution of HIV-1 promoter methylation *ex vivo* to EGCG reactivation potency. A dynamical increase in the degree of viral promoter methylation has been shown in HIV^+^ individuals in response to the duration of the antiretroviral treatment or the time of viral suppression (16, 17). Accordingly, strong statistically significant positive correlations were observed between the EGCG-mediated HIV-1 reactivation potency and either time on cART (Fig. S9A) or time of virological suppression (Fig. S9B), while no correlation was observed for the positive control (Fig. S9C). These results demonstrate that EGCG modes of action *ex vivo* are time-dependent in HIV^+^ individuals, suggesting that EGCG acts, at least in part, through HIV-1 promoter demethylation. To refine this observation, we next assessed the DNA methylation level of the viral promoter in our cohort of HIV^+^ individuals by sodium bisulfite sequencing. As suggested by Blazkova and colleagues (10), we found that the assessment of DNA methylation in aviremic individuals, who have smaller reservoirs that prevent the analysis of proviral DNA, was technically challenging. Nevertheless, we obtained the DNA methylation profile of the HIV-1 promoter for 8 out of the 22 enrolled HIV+ individuals. Out of these, five individuals had no detectable DNA methylation, while three individuals presented a median methylation level of 7.41% mCpG on the HIV-1 promoter, which corresponds to levels reported in other studies (10, 12, 17). To determine the relationship between EGCG reactivation potency and *in vivo* HIV- 1 promoter methylation, we clustered these 8 individuals in groups of non-methylated or methylated 5’LTR and plotted the cell-associated HIV-1 US RNA for 50µg/mL of EGCG (Figure S9D). Without reaching statistical significance, a trend towards a higher reactivation could be observed for patients accumulating more DNA methylation on the viral promoter. The lack of statistical significance can be attributed to the low number of patients but it also reveals the pleiotropic reactivation capacity of EGCG. We propose that the mechanism of EGCG reactivation of HIV-1 from latency is through DNA demethylation of the viral promoter, or, when it is not methylated, through indirect demethylation of cellular genes.

Altogether, our *ex vivo* data indicate, in line with the heterogeneity of the mechanisms responsible for HIV-1 latency, that EGCG, in addition to its antiviral properties, is a heterogeneous LRA, capable of reversing HIV-1 latency through several modes of actions depending on different infected individuals.

## DISCUSSION

Accumulating data highlights the intrinsically dynamic and heterogeneous nature of latent HIV- 1 cellular reservoirs within and between infected individuals. This heterogeneity and the multiplicity of the silencing mechanisms underlying HIV-1 latency rather than latency in itself are now considered as the major barrier to eradicating HIV-1 (8). In agreement, LRAs have been found to present various reactivation potencies *in vitro* and *ex vivo* (45–47). In the context of DNA methylation, our previous study has highlighted that the DNA methylation inhibitor 5-AzadC exhibits different *ex vivo* reactivation potencies in terms of HIV-1 latency reversal (18). Here, we investigated the molecular source of this potency heterogeneity at the level of proviral DNA methylation.

We first evidenced the existence of non-random and reproducible DNA methylation signatures in response to 5-AzadC treatment at the level of the single CpG dinucleotide in the HIV-1 promoter. Thus, rather than reactivating HIV-1 in a non-specific manner, 5-AzadC acts through specific molecular mechanisms within the viral promoter. To tease out for these regulatory mechanisms, we mapped preferentially-demethylated positions in the 5’LTR. In the present study, we focused on the most significantly-demethylated CpG position of our whole dataset but a similar approach can be used for other positions or for other epigenetic LRAs. DDMP5 is located in a CRE, however, rather than showing that DDMP5 methylation prevents the binding of cognate transcriptional activators (22, 48), we demonstrated that DDMP5 methylation allowed the *in vitro* binding of an additional protein complex, containing UHRF1. We also reported *in vivo* recruitment of UHRF1 to the latent HIV-1 promoter. Interestingly, we showed that UHRF1 recruitment to the latent promoter was only proportional to the level of DDMP5 methylation, indicating that UHRF1 could be recruited redundantly to the latent promoter through other mechanisms than DNA methylation. This redundancy is a general trend in the recruitment of epigenetic machineries to the HIV-1 5’LTR and safeguarding their recruitment points towards the importance of the epigenetic repression of viral genes during latency (8). UHRF1, in particular, is an important epigenetic integrator as it both reads the chromatin (5mC, H3K9me3 and H3R2) and recruits epigenetic enzymes (DNMT1, G9a/EHMT2, …). We thus investigated the role of the recruited UHRF1 in participating in the 5’LTR heterochromatinization during HIV-1 latency. We report here for the first time a role for UHRF1 in HIV-1 silencing through DNA methylation. Mechanistically, UHRF1 knockdown led to a statistically significant decrease in the global DNA methylation level of the 5’LTR and of DNMT1 recruitment. We also showed that G9a/EHMT2 recruitment to the 5’LTR was decreased upon UHRF1 knockdown, supporting the model that UHRF1 branches several repressive epigenetic mechanisms during HIV-1 latency. Considering the multiple modes of gene repression mediated by UHRF1, we further sought to determine how much of its repressive action on the 5’LTR was dependent on DNA methylation, and in particular on the methylation status of DDMP5. To answer unambiguously to this question, we decided to work in an artificial system of transient transfection and showed that, even there, UHRF1 repression of HIV-1 transcription depended strongly on DNA methylation and, in a large part, on DDMP5. Thus, in agreement with its potential multiple recruitment modes to the viral promoter, UHRF1 represses HIV-1 gene expression during latency through several mechanisms, but predominantly, through DNA methylation. Together, these results demonstrate that UHRF1 is a new actor in HIV-1 latency.

As a proof-of-concept that the molecular characterization of HIV-1 latency leads to the identification of new targets for therapeutic strategies, we next studied the latency-reversing potency of UHRF1 downregulation. EGCG, the major phenolic compound of green tea, has been reported, among other functions, to affect UHRF1 expression (49). Using complementary models – latently-infected J-Lat T-cell lines and *ex vivo* cell cultures from cART-treated HIV^+^ aviremic individuals – our results showed the potency of EGCG as a new LRA. Indeed, EGCG reactivated HIV-1 from latency up to the completion of the viral replication cycle, in a short time frame, and in all the latency models we tested. Interestingly, EGCG had been reported to present an antiviral activity on HIV-1, by inhibiting several steps of its replication cycle, notably, by binding and destabilizing the HIV-1 envelope (38). This has led to its use in anti-HIV clinical trials (NCT01433289 and NCT03141918, ClinicalTrials.gov). In agreement with EGCG antiviral activity on HIV-1 (38), we detected little p24^Gag^ capsid protein in the cell culture supernatants in our reactivation experiments. We nevertheless observed, in our *ex vivo* patient cell cultures, that EGCG augmented the recovery of extracellular HIV-1 RNA. Thus, these results suggest that reactivated virions were indeed destabilized by EGCG in cell culture supernatants. Furthermore, we showed that EGCG treatment was associated with a decreased expression of the cellular surface marker CD4 on target cells, thereby preventing subsequent new infections following reactivation. We thus showed the promising use of EGCG in HIV-1 eradication strategies due to its complementary and synergistic anti-HIV modes of action. On a larger perspective, we also showed the relevance of targeting UHRF1 in HIV-1 cure therapeutic strategies, in trend with the considerable attention for UHRF1 inhibitors in the cancer field (49). Our results indicate, however, that EGCG potency is not uniquely dependent on UHRF1 expression, rather this compound presents a pleiotropic function in HIV-1 reactivation, balancing the heterogeneity observed in viral reservoirs.

In conclusion, we developed a probabilistic methodology to decipher the DNA methylation-mediated mechanisms underlying the heterogeneous capacity of 5-AzadC to reactivate HIV-1 from latency. This approach enabled us to uncover the role of UHRF1, an epigenetic integrator, in HIV-1 promoter silencing *via* DNA methylation. Our work provides a demonstration that the understanding of the molecular basis of the heterogeneous effects of LRAs, even in *in vitro* HIV-1 latency models, can bring to light new factors involved in HIV-1 silencing and hence, new targets to devise anti-HIV therapeutic approaches. As a proof-of-concept, we showed that EGCG, a known inhibitor of UHRF1, presents, in addition to its broad antiviral activities, a pleiotropic anti-latency activity, allowing its potential use in HIV-1 cure strategies.

## ACKNOWLEDGMENTS

We thank the members of the ANRS (French National Agency for Research on AIDS and Viral Hepatitis) RHIVIERA (Remission of HIV Era) Consortium for helpful discussions. We thank the HIV- 1^+^ individuals for their willingness to participate in this study. We thank the nursing team of CHU Saint-Pierre Hospital (Elodie Goudeseune, Joëlle Cailleau and Annick Caestecker) who cared for the HIV^+^ individuals. We thank Jacqueline Pineau from the transfusion center of Charleroi (Belgium) for providing blood from healthy donors. We thank Hilde Vereertbrugghen from Francis Corazza’s laboratory for excellent technical assistance. We thank Motoko Unoki for precious advice regarding UHRF1 RNA interference. We thank Mitia Duerinckx and Benoît Van Driessche for probabilistic and statistics advice.

CVL acknowledges funding from the Belgian National Fund for Scientific Research (FRS-FNRS, Belgium), the « Fondation Roi Baudouin », the NEAT (European AIDS Treatment Network) program, the Internationale Brachet Stiftung, ViiV Healthcare, the Walloon Region (« Fonds de Maturation »), « Les Amis des Instituts Pasteur à Bruxelles, asbl », and the University of Brussels (Action de Recherche Concertée ULB grant) related to her work on HIV latency. The laboratory of CVL is part of the ULB-Cancer Research Centre (U-CRC). RV was funded by an “Aspirant” fellowship (F.R.S-FNRS) and a fellowship from “Les Amis des Instituts Pasteur à Bruxelles, asbl” and is a Belgian American Educational Foundation (BAEF) fellow and a scientific collaborator of the ULB. LN is supported by a “PDR” grant from the F.R.S-FNRS. GD is postdoctoral clinical master specialist for the F.R.S-FNRS. A A-A is a fellow of the “Wallonie-Bruxelles International” Program and of the Marie Skłodowska Curie COFUND action. EP is a fellow of the “Télévie Program” (F.R.S-FNRS). MB is funded by a FRIA fellowship (F.R.S.-FNRS). CVL is “Directeur de Recherches” at the F.R.S-FNRS. Work in OR’s laboratory was supported by grants from the French agency for research on AIDS and viral hepatitis (ANRS), Sidaction and Alsace contre le Cancer. This project has received funding from the European Union’s Horizon 2020 research and innovation program under grant agreement No 691119- EU4HIVCURE-H2020-MSCA-RISE-2015.

## AUTHOR CONTRIBUTIONS

Conceived and designed the experiments: CVL, RV and SB. Performed the experiments: RV, SB, LN, GD, CV, AA-A, MB, EP, ND. Performed the HIV-1 DNA and RNA quantifications: AP, GD, VAF and AC. Performed the activation analyses: FC. Performed the primary models of HIV latency experiments: VLD, VG. Performed the Infinium analyses: MB and FF. Performed the ultra-sensitive p24 assays: AS-C. Performed HIV^+^ individuals selection: CN, SDW. Analyzed the data: RV, SB, LN, GD, CV, AA-A, MB, EP, ND and CVL. Contributed in reagents/materials/analysis tools: CS, CR, BB, VG, OR. Wrote the paper: RV, SB and CVL.

## MATERIAL AND METHODS

### Cell culture

Jurkat, J-Lat 6.3, J-Lat 8.4, J-Lat 9.2, J-Lat 15.4 and the HEK293T cell lines were obtained from the AIDS Research and Reference Reagent Program (NIAID, NIH). Cells were grown in RPMI 1640 medium (Gibco-BRL) supplemented with 10 % fetal bovine serum, 50 U/ml of penicillin and 50 µg/mL of streptomycin and were cultivated at 37°C in a 5% CO2 atmosphere.

### Reagents and antibodies

5-aza-2’-deoxycytidine (5-AzadC, A3656) and epigallocatechin-3-gallate (EGCG, E4143) were purchased from Sigma Aldrich. Antibodies against CREB (sc-186), CREM (sc-440), ATF-1 (sc-28673), MBD4 (sc-10753) and purified rabbit IgG (sc-2027) were purchased from Santa Cruz Biochemical. Antibodies against MBD1 (pAb-078-050) and UHRF1 (H00029128-B01P) were purchased from Diagenode and Abnova, respectively. Antibodies against MBD2/3 (07-199) and Kaiso (05-659) were purchased from Upstate/Millipore. Antibodies against MeCP2 (ab2828) and RBP-JK (ab33065) were purchased from Abcam. Antibodies against RNAPII (14958) were purchased from Cell Signaling. Secondary antibodies were purchased from Cell Signaling (7074 and 7076).

### Virus production assays

HIV-1 production was measured in cell culture supernatants of cell cultures by ELISA assays on p24^Gag^ using the INNOTEST HIV Antigen mAb kit per the manufacturer’s instructions (Fujirebio).

### Sodium Bisulfite-mediated mapping of methylcytosines

Genomic DNA was isolated using the DNeasy Blood and Tissue kit (Qiagen), then sodium bisulfite-converted (EpiTect Bisulfite kit, Qiagen). The 5’LTR or the *rev* regions were amplified by (semi)nested PCR (primer sequences are available upon request). At least 12 clones from each condition were sequenced, and clones with sodium bisulfite conversion higher than 95% were aligned on the HIV- 1 NL4.3 reference sequence using ClustalΩ. MethTools (50) and Inkscape were used for graphical representations.

### Electrophoretic Mobility Shift Assays (EMSAs)

Nuclear extracts were prepared using a protocol described by Dignam and colleagues (51).

EMSAs, competition EMSAs and supershift assays were performed as described previously (48).

### Chromatin Immunoprecipitation assays

ChIP assays were performed as previously described (52), using the ChIP assay kit from Millipore or equivalent homemade solutions. Relative quantification using standard curve method on the input was performed for each primer pair and 96-well Optical Reaction plates were read in a StepOnePlus PCR instrument (Applied Biosystems). Fold enrichments were calculated as fold inductions relative to the values measured with IgG. Primer sequences used for quantification are available upon request.

### Western Blot

Western blotting was performed with 15μg of total protein extracts. The immunodetection was assessed using primary antibodies targeting UHRF1 or β-actin as loading control. Horseradish peroxidase (HRP)-conjugated secondary antibodies were used for chemiluminescence detection (Cell Signaling).

### RNA extraction and analysis of transcripts

Total RNA samples were isolated using the Tri-Reagent (TRC-118, MRC), according to the manufacturer’s protocol. Following DNAse treatment (AM1907, Invitrogen), reverse transcription was performed with the PrimeScript RT reagent kit (RR037A, TaKaRa).

### Lentiviral production and transduction assays

TRC Lentiviral shRNA plasmids (pLKO.1) MISSION shRNA were obtained from Sigma-Aldrich (SHC002, TRCN0000273315, TRCN0000273256, TRCN0000273317 and TRC0000004352).

The pMD2.G and the psPAX2 packaging system were obtained from Addgene. VSV-G pseudotyped particles were produced by transfection of HEK 293T cells as described previously(33). J-Lat cells were transduced as described previously (53).

### Cellular proliferation assays and viability

Cellular proliferation was evaluated by the colorimetric test WST-1 according to the manufacturer’s instructions (Roche). Cellular viability was assessed by staining the cells with the LIVE/DEAD Fixable Near-IR Dead Cell Stain (Thermo Fisher) and analysis by flow cytometry on a FACSCantoII (Becton-Dickinson), using the FACSDiva Software (Becton-Dickinson).

### Plasmid constructs and reporter assays

The human UHRF1 expression vector (pCMV-HA-UHRF1, termed pUHRF1) was kindly provided by Dr Olivier Rohr. The non-episomal pLTR-Fluc vector was described previously (54). To obtain the pLTRme-Fluc vector, where only the LTR CpGs are methylated, we methylated *in vitro* the whole pLTR-Fluc construct using the SssI methyltransferase (New England Biolabs, M0226). The LTR fragment was then purified, cloned back in the parental reporter vector and the resulting pLTRme-Fluc vector was directly transfected without bacterial amplification.

### Infinium 850K Human Methylation arrays

Genomic DNA was extracted with the DNeasy Blood and Tissue Kit (QIAGEN) and converted with sodium bisulfite (EZ DNA Methylation Kit, Zymo Research). The quality of each analyzed sample was first evaluated by inspection of the control probes intensity level. Raw data (uncorrected probe intensity values) from the Infinium Methylation arrays were processed according to the recommended steps of Dedeurwaerder *et al.*(55). Beta-values were computed using the following formula: Beta-value = M/[U+M] where M and U are the raw “methylated” and “unmethylated” signals, respectively. Beta-values were corrected for type I and type II bias using the peak-based correction (56). Infinium HumanMethylation850K raw data were submitted to the NCBI’s Gene Expression Omnibus (GEO) database (GSE139320, token : wnqfssiudxubtgx). Principal component analysis and hierarchical clustering were performed via an in-house R script using the most variable Infinium probes (standard deviation ≥ 0.27). For differential analysis, probes showing an absolute difference between case and control Beta-values higher than 0.3 were assumed significant. For pathway analysis, Infinium probes located in promoter regions were first associated to their corresponding genes. Then, a « delta-Beta » was defined for each gene as the difference between case and control Beta-values of its promoter Infinium probe showing the highest absolute difference. Finally, genes were ranked according to their delta-Beta and submitted to the GSEA tool (41) to search for significant enrichments among the HALLMARK gene sets from MSigDB (http://software.broadinstitute.org/gsea/msigdb/).

### Study subjects

We selected 22 HIV-1-infected individuals at the Saint-Pierre Hospital (Brussels, Belgium) based on the following criteria: all volunteers were treated with cART for at least 1 year, had an undetectable plasma HIV-1 RNA level (20 copies/ml) for at least 1 year, and had a level of CD4+ T lymphocytes higher than 300 cells/mm^3^ of blood. Characteristics (age, CD4+ T cell count, CD4+ nadir, antiviral regimens, duration of therapy, duration with undetectable plasma HIV-1 RNA level, and HIV-1 subtypes) of HIV^+^ individuals from the Saint-Pierre Hospital were well documented and are presented in Table S1A. Buffy coats from healthy donors were obtained at the Belgian Red Cross.

### Ethical statement

Ethical approval was granted by the Human Subject Ethics Committee of the Saint-Pierre Hospital (Brussels, Belgium). All individuals enrolled in the study provided written informed consent for donating blood.

### Isolation of CD8^+^-depleted PBMCs

CD8^+^-depleted PBMCs used in reactivation assays were isolated from fresh whole blood of HIV^+^ individuals as previously described (18).

### Quantitation of cell-associated HIV-1 unspliced RNA

Total nucleic acids were extracted from pellets of CD8^+^-depleted PBMCs according to the Boom isolation method (57). Extracted cellular RNA was treated with DNase (DNA-free kit; Thermo Fisher Scientific) and reverse transcribed using the SuperScript III reverse transcriptase (Thermo Fisher Scientific). cDNA was used for the qPCR-based quantification of cell-associated HIV-1 unspliced RNA (amplicon in the *gag* region), as previously reported (58). HIV-1 RNA copy numbers were normalized to the total cellular RNA (by measurement of 18S ribosomal RNA) inputs as described previously (59). Non-template control wells were included in every qPCR run and were consistently negative.

### Quantification of HIV-1 extracellular RNA

Total RNA was extracted from CD8^+^-depleted PBMCs *ex vivo* culture supernatants using the QIA amp Viral RNA Mini kit (Qiagen). HIV-1 RNA levels were quantified using the Generic HIV Viral Charge kit (Biocentric) according to the manufacturer’s instructions.

### Quantification of total HIV-1 DNA

Total cellular DNA was extracted from patient CD8^+^-depleted PBMCs *ex vivo* cultures using the QIAamp DNA Mini kit (Qiagen). The total cell-associated HIV-1 DNA was then quantified by ultra- sensitive real-time PCR (Generic HIV DNA cell kit, Biocentric) according to the manufacturer’s instruction (60).

### Cell activation analysis by flow cytometry

For cell activation analysis, CD8^+^-depleted PBMCs from blood of healthy donors were used to establish *ex vivo* cell cultures. Cells were collected 24 hours after stimulation with EGCG and were stained with relevant antibodies as previously described (18).

### Statistical analysis

The demethylation probability following 5-AzadC treatment was calculated as follows:

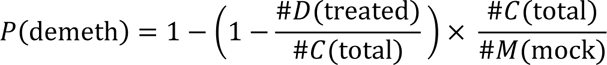

where P(demeth) corresponds to the probability of demethylation following treatment, #D(treated) the number of demethylated CpGs in the treated conditions, #C(total) the number of clones (12 for each condition in the present study) and where #M(mock) corresponds to the number of methylated CpGs in the mock-treated conditions. Sodium bisulfite sequencing data sets were analyzed using Fisher’s exact test. For all the analyses, the threshold of statistical significance was set at 0.05. p-values ≤ 0.05 (*: p- value ≤ 0.05, **: p-value ≤ 0.01, ***: p-value ≤ 0.001) were considered statistically significant. All tests were two-sided. All analyses were performed using Prism version 6.0 (GraphPad software) and Microsoft Excel. Statistical tests are indicated in the corresponding figure legends.

---

**Figure S1:**
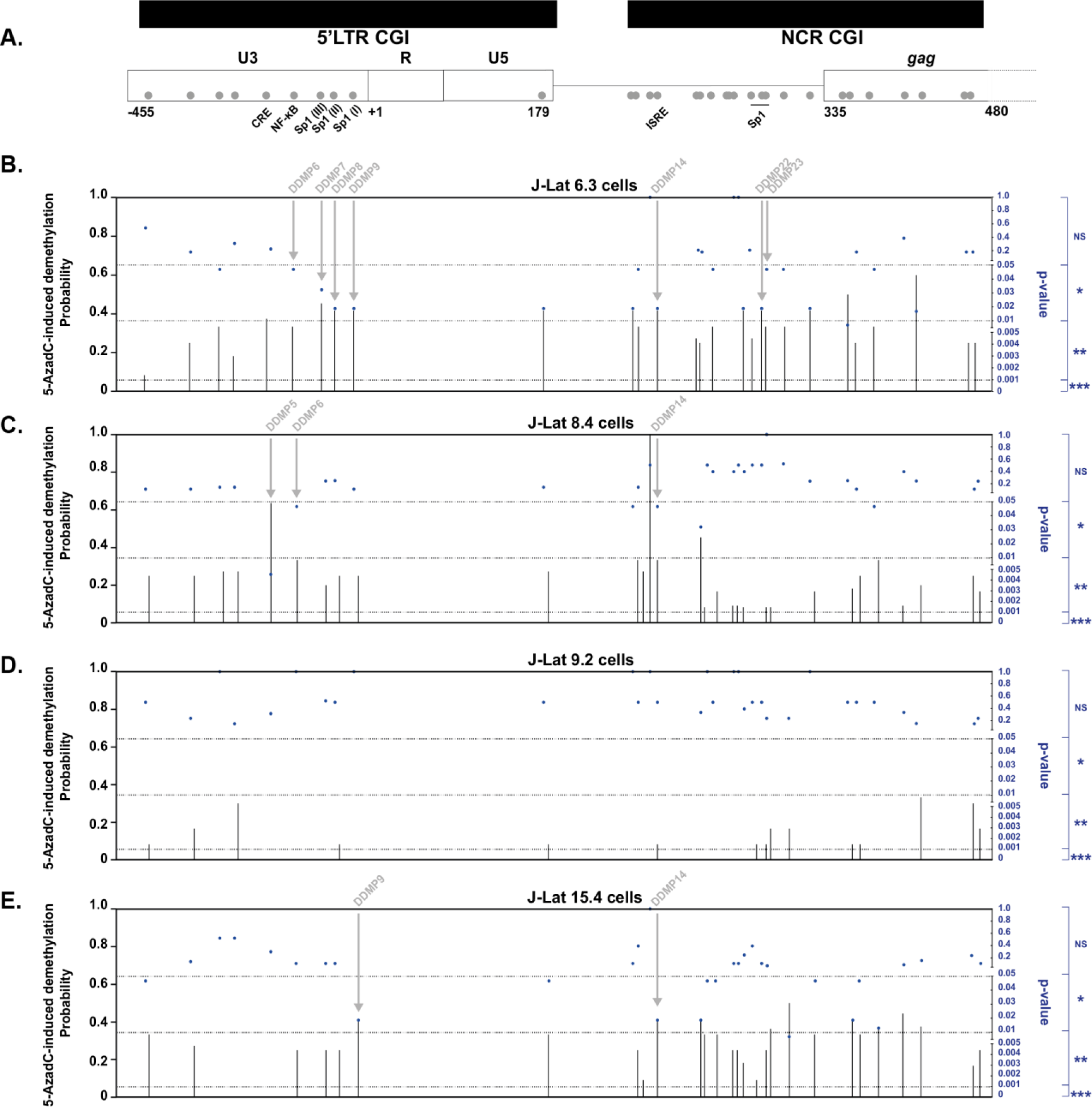
Mapping of 5-AzadC-induced demethylation probability reveals hotspots of demethylation. **(A)** Schematic view of CpG dinucleotides position in the 5’LTR and transcription factor binding sites positions. The individual probabilities of 5-AzadC-induced demethylation at each CpG dinucleotide, presented in histograms on the left Y axis and associated p-values (Fisher’s exact test), presented in blue on the right Y axis, was calculated in J-Lat 6.3 cells **(B)**, in J-Lat 8.4 cells **(C)**, in J-Lat 9.2 cells **(D)** and in J-Lat 15.4 cells **(E)** using the data presented in Figure 1. Statistically significant DDMPs located in transcription factor binding sites are highlighted and indicated by an arrow.

**Figure S2:**
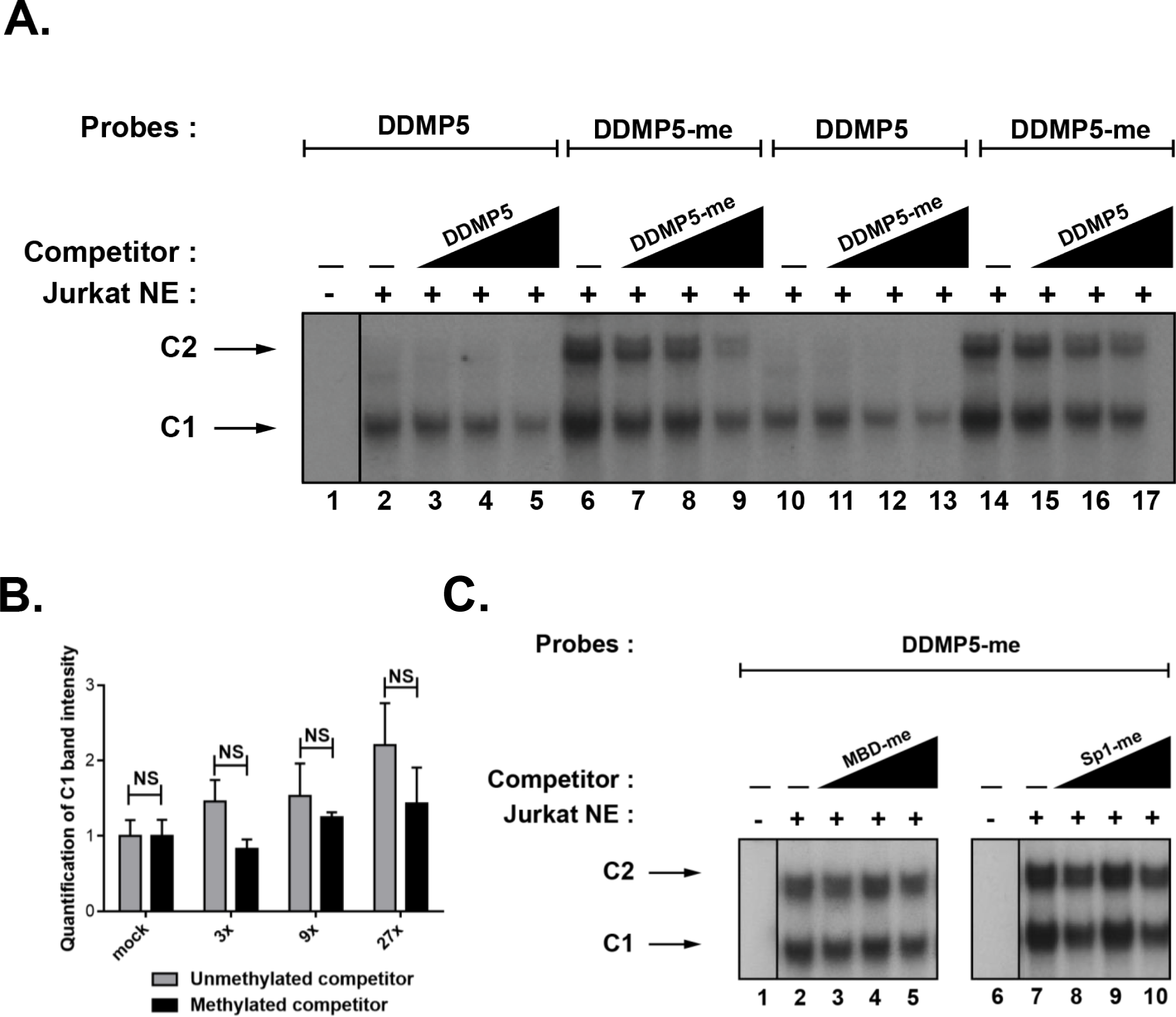
Methylation of DDMP5 promotes the sequence-specific and methylation-specific *in vitro* binding of an additional factor. **(A)** The radiolabeled unmethylated or methylated HIV-1 DDMP5 probe was incubated with 10µg of Jurkat cells NE, in the absence of competitor, or in presence of increasing molar excesses of the unlabeled homologous, respectively methylated (lanes 7-9 and lanes 11-13) or unmethylated (lanes 3-5 and lanes 15-17), HIV-1 DDMP5 oligonucleotides. **(B)** The quantification of the band corresponding to the C1 complex for the competition with the unmethylated competitor (equivalent to the lanes 3-5) and the methylated competitor (equivalent to the lanes 7-9) is presented. Statistical significance was calculated by an unpaired T test. **(C)** The methylated HIV-1 DDMP5 radiolabeled probe was incubated with 10µg of Jurkat NE, in the absence of competitor, or in presence of increasing molar excesses of the heterologous unlabeled methylated MBD consensus (indicated as “MBD-me”, lanes 3-5)) or methylated Sp1 consensus (indicated as “Sp1-me”, lanes 7-10) oligonucleotides. Binding reactions were analyzed by PAGE, and retarded complexes were visualized by autoradiography. The major DNA-protein complexes C1 and C2 are indicated by arrows. The figure shows only the specific retarded bands of interest. One representative experiment out of three is presented.

**Figure S3:**
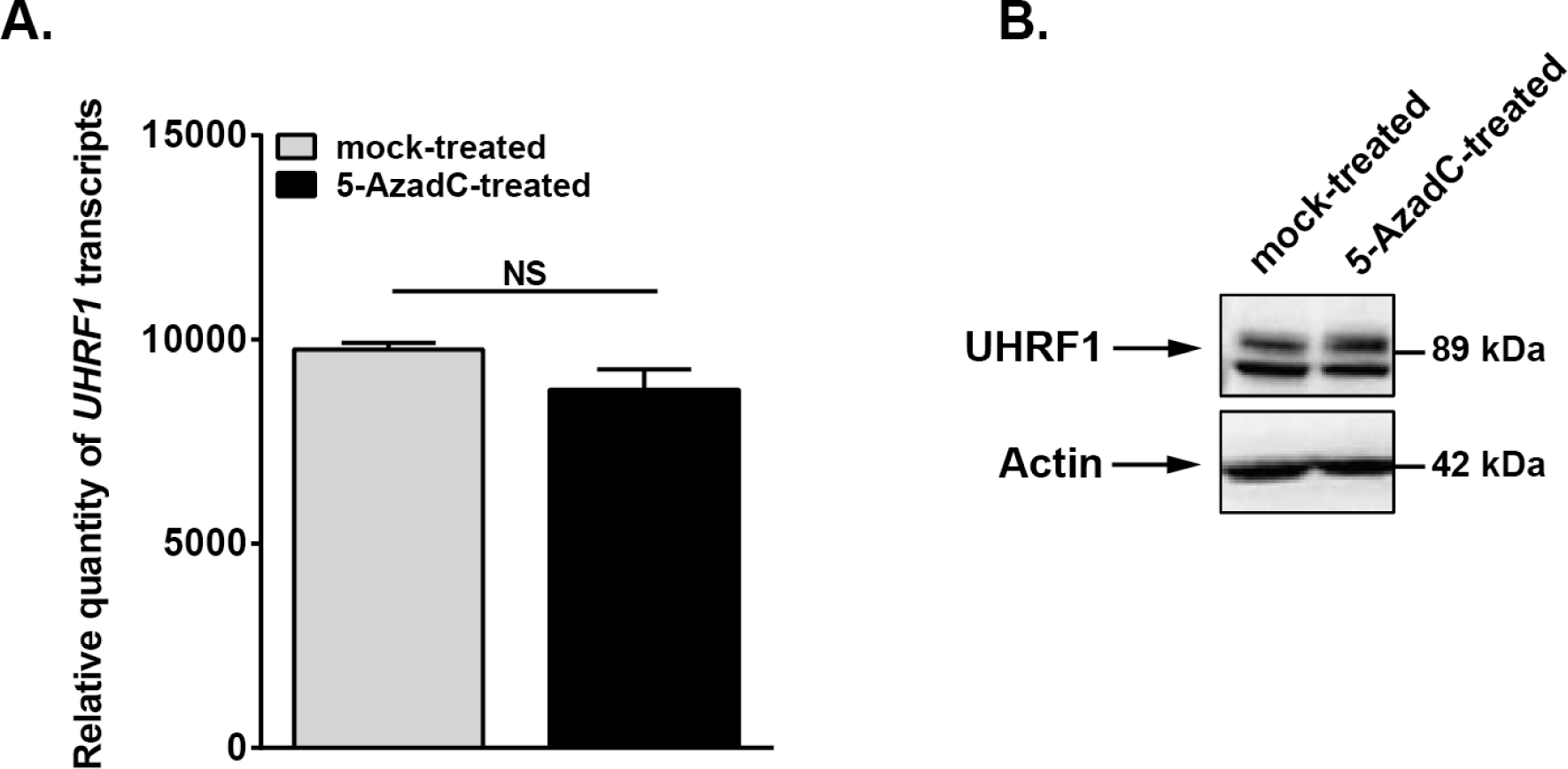
UHRF1 expression is not affected by 5-AzadC. **(A)** Total RNA preparations from the J-Lat 8.4 cells, either mock-treated or treated with 400nM of 5-AzadC for 72h were reverse transcribed. UHRF1 transcripts were quantified by reverse transcription qPCR using GAPDH as a normalizer. Means from duplicates ± SD are indicated. Statistical significance was calculated with an unpaired T test. One representative experiment out of three is presented. **(B)** Total protein extracts from the same stimulation as in (D) were extracted. UHRF1 was immunodetected by western blotting using a specific antibody. Levels of β-actin were measured to control protein loading. One representative experiment out of three is presented.

**Figure S4:**
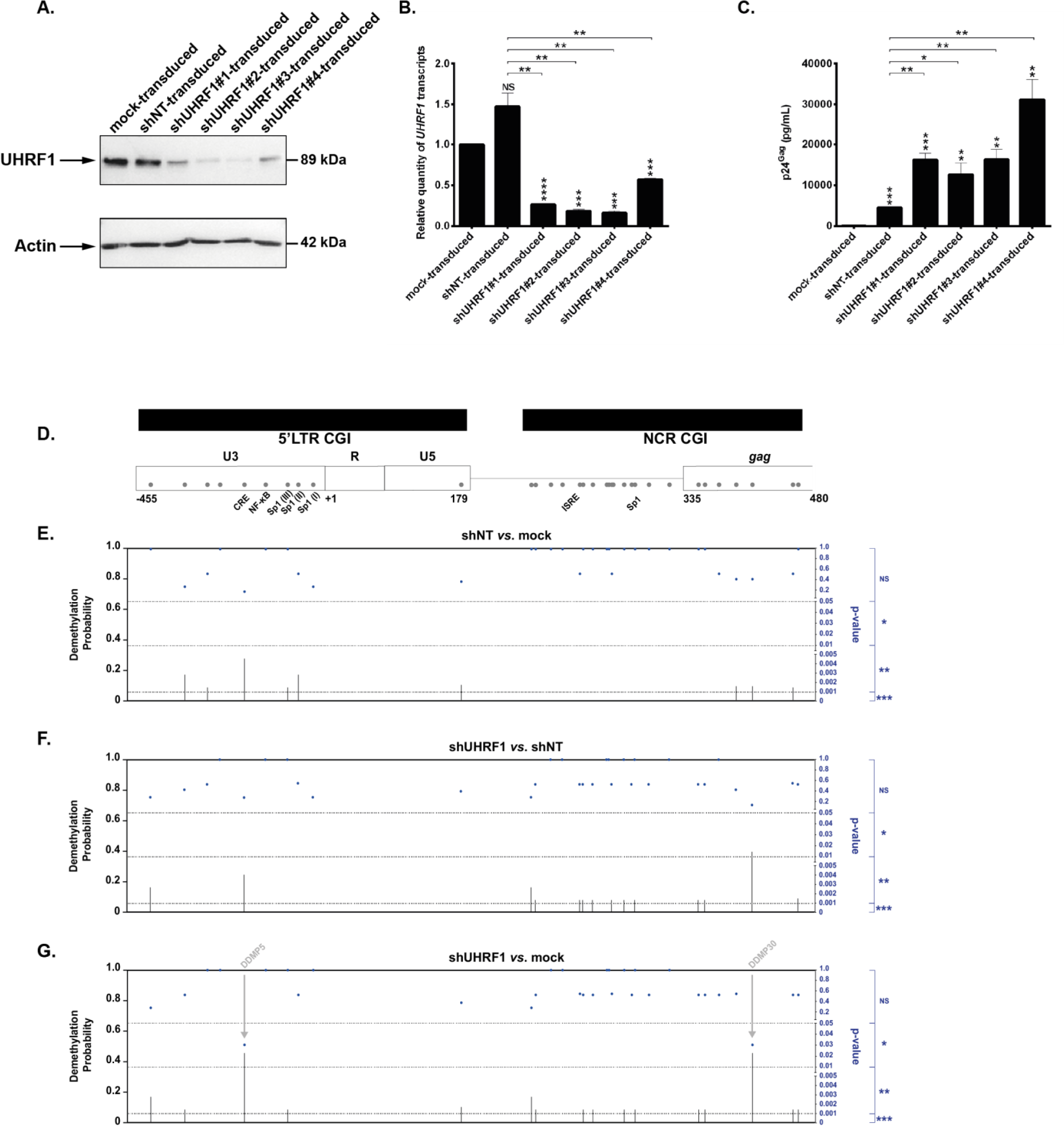
UHRF1 depletion in J-Lat 8.4 cells following shUHRF1 transduction provokes reactivation of HIV-1 and specific demethylation patterns in the viral promoter. **(A)** UHRF1 and β-actin, serving as a loading control, protein levels were assessed by immunoblot in whole-cell lysates of J-Lat 8.4 cells mock-transduced, transduced with a control shRNA (indicated as “shNT-transduced”) or transduced with four independent shUHRF1 (indicated as “shUHRF1#1-4- transduced”). **(B)** Total RNA preparations were used in RT-qPCR to quantify UHRF1 transcripts, using GAPDH as a first normalizer and the mock-transduced condition as second normalizer. Values correspond to means ± SD of two independent qPCRs. **(C)** The quantity of p24^Gag^ in the culture supernatants was evaluated by ELISA. Results are representative of the mean ± SD two independent ELISA quantifications. The whole panel originates from one representative experiment out of three. Statistical significance was assessed by an unpaired T test and corresponds, if not otherwise specified, to comparisons to the mock-transduced condition. **(D)** Schematic view of CpG dinucleotides position in 5’LTR and transcription factor binding sites positions. The individual probabilities of 5-AzadC-induced demethylation at each CpG dinucleotide, presented in histograms on the left Y axis and associated p-values (Fisher’s exact test), presented in blue on the right Y axis, was calculated in J-Lat 8.4 cells shNT-transduced in comparison to mock-transduced cells **(E)**, in J-Lat 8.4 cells shUHRF1-transduced in comparison to shNT-transduced cells **(F)** and in in J-Lat 8.4 cells shUHRF1-transduced in comparison to mock-transduced cells **(G)**, using the data presented in Figure 3. Statistically significant DDMPs located in transcription factor binding sites are highlighted.

**Figure S5:**
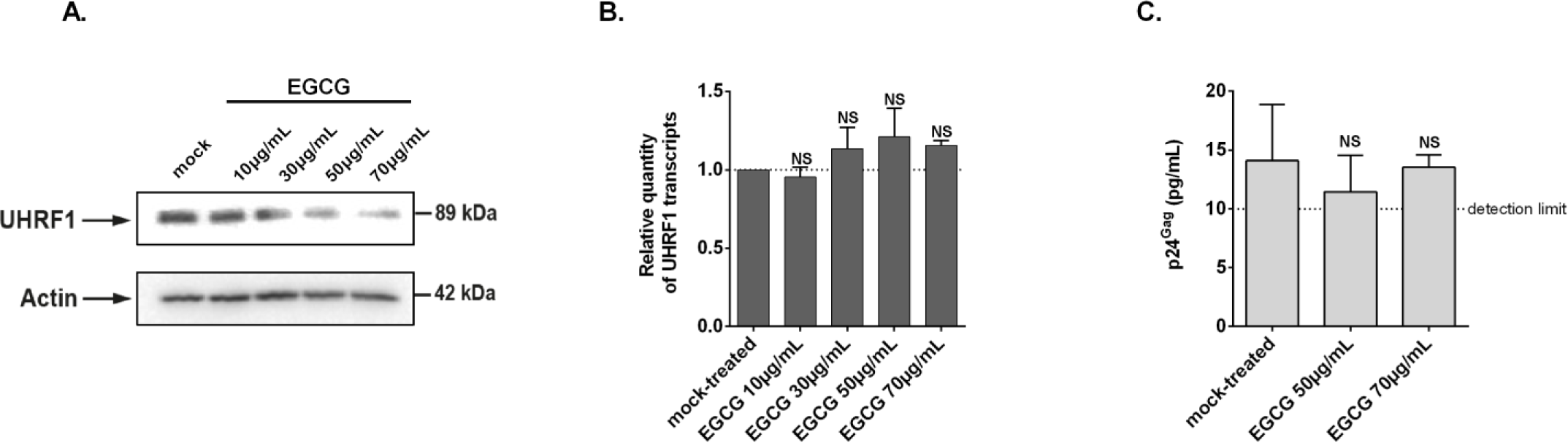
UHRF1 is downregulated in J-Lat 8.4 cells following EGCG treatment. **(A)** UHRF1 and β-actin, serving as a loading control, protein levels were assessed by immunoblot in whole-cell lysates of J-Lat 8.4 cells mock-treated or treated with increasing doses of EGCG for 24h. **(B)** Total RNA preparations from the same experiment as in A were used in RT-qPCR to quantify UHRF1 transcripts, using GAPDH as a first normalizer and the mock-treated condition as second normalizer. One representative experiment is shown out of three. Statistical significance was assessed using an unpaired T test. **(C)** The quantity of p24^Gag^ in the culture supernatants of mock-treated or EGCG-treated J-Lat 8.4 cells was evaluated by ELISA. Results are representative of the mean ± SD of three independent EGCG treatment. The limit of the test detection at 10pg/mL is indicated. Statistical significance was assessed using an unpaired T test.

**Figure S6:**
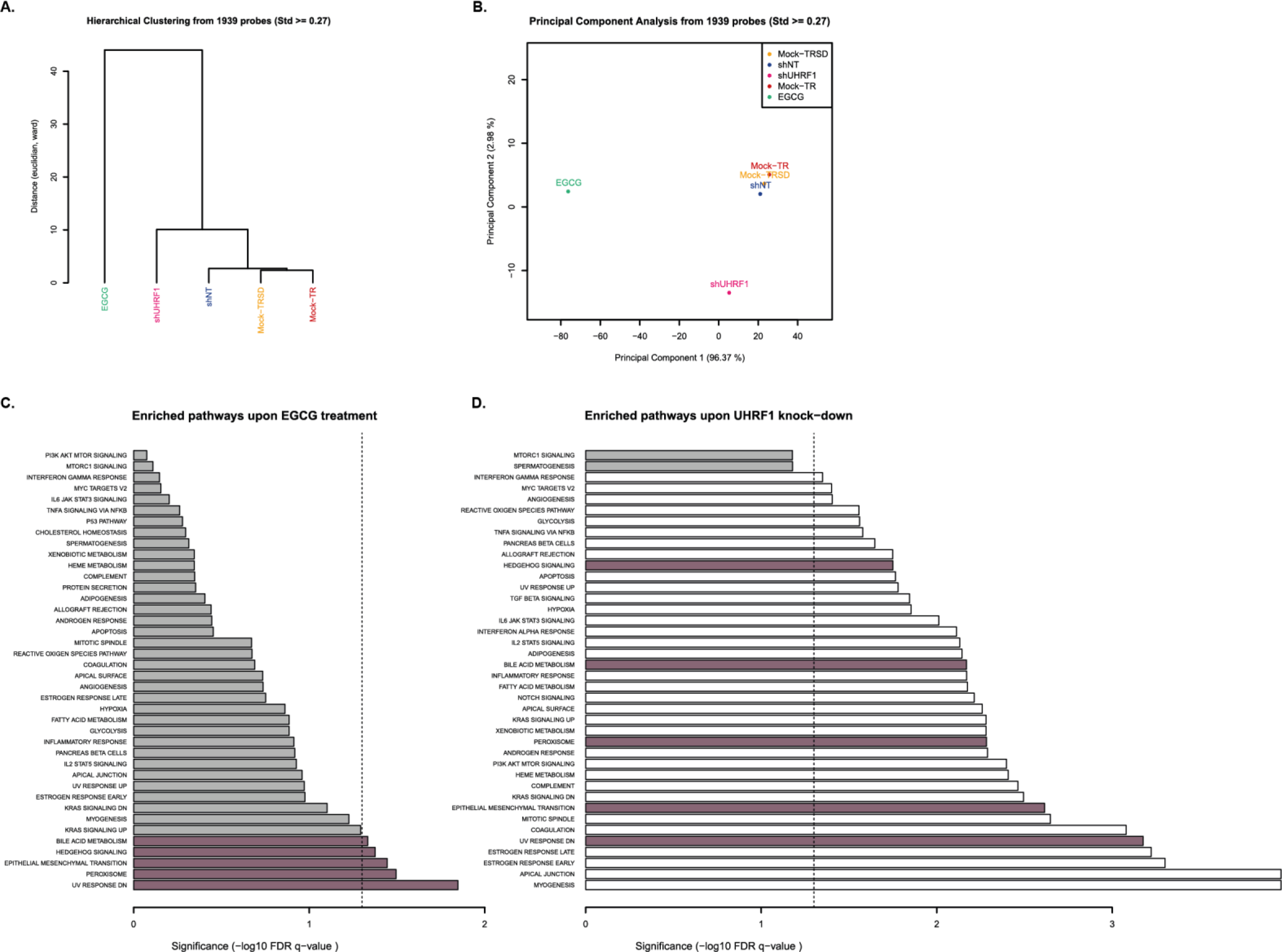
Methylome-wide analyses of shUHRF1 and EGCG-induced hypomethylation show different patterns. **(A)** Hierarchical clustering was performed on the most variable CpGs obtained in the Infinium assay for the genomic DNA samples of Fig. 3 and Fig. 4. **(B)** Principal component analysis was performed on the most variable CpGs obtained in the Infinium assay for the genomic DNA samples of Fig. 3 and Fig. 4. **(C)** Enriched pathways corresponding to the differential DNA methylation profile upon EGCG treatment were analyzed by GSEA. Grey bars represent non-significant pathways, purple bars represent statistically significative common pathways and white bars represent statistically significative but not common pathways. **(D)** Enriched pathways corresponding to the differential DNA methylation profile upon shUHRF1 transduction were analyzed by GSEA.

**Figure S7:**
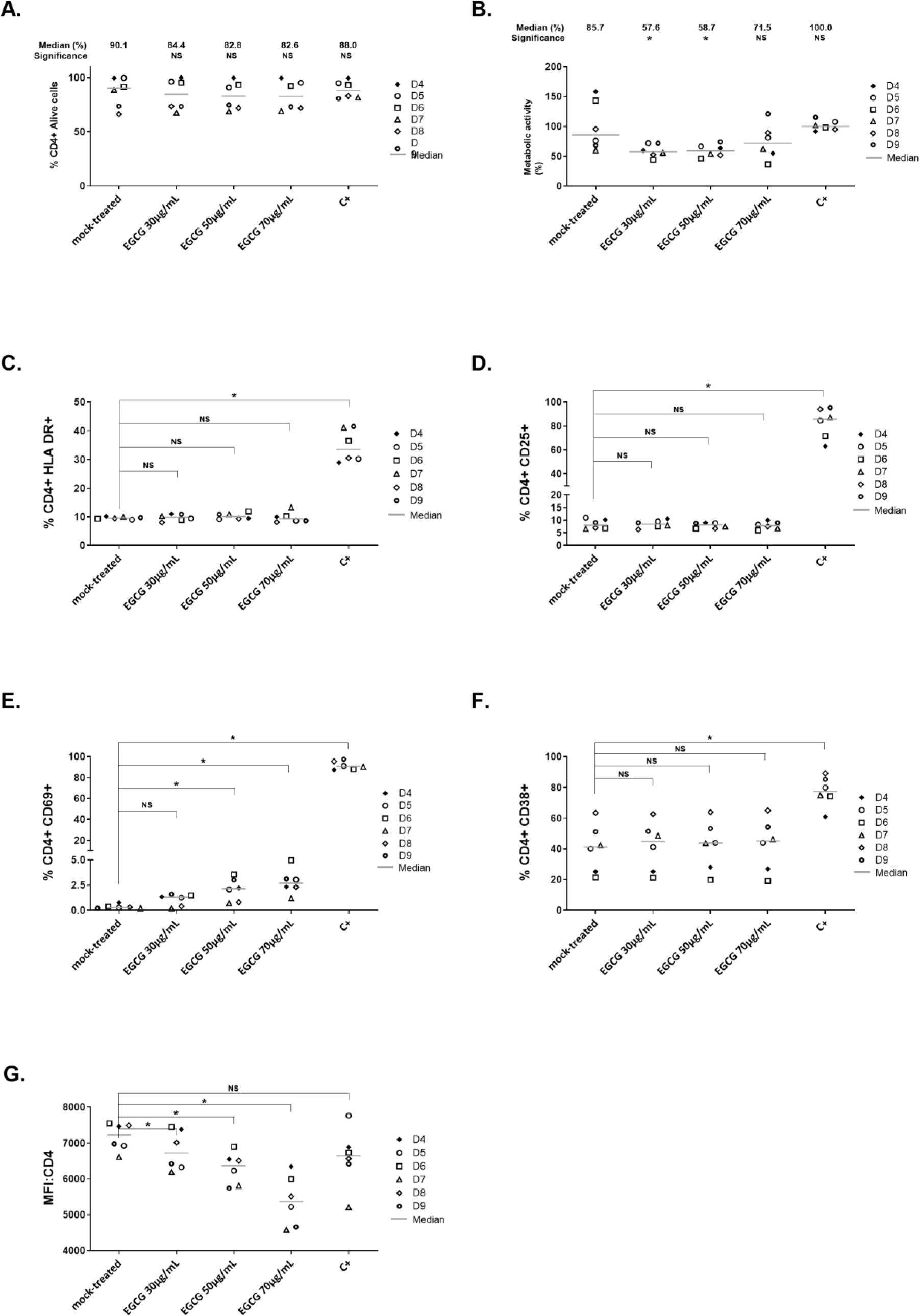
Effects of EGCG on cellular viability, metabolic activity, cell surface activation markers and CD4 expression in CD8^+^-depleted PBMCs from healthy donors. CD8^+^-depleted PBMCs were extracted from six healthy donors blood samples and were mock-treated or treated for 24h with the indicated doses of EGCG, or with anti-CD3+anti-CD28 antibodies serving as positive control stimulation. **(A)** Cells were stained for CD4+ and LIVE/DEAD Fixable Near-IR Dead Cell Stain to discriminate between viable and non-viable cells. Medians percentages are indicated. **(B)** WST-1 proliferation assay, reflective of metabolic activity, was performed on the cells. Median percentages are indicated. HLA-DR **(C)**, CD25 **(D)**, CD69 **(E)** and CD38 **(F)** expression was analyzed by flow cytometry. Results are presented as percentage of marker expression in alive CD4+- gated cell populations. Medians for each condition are presented. **(G)** Mean fluorescence intensify (MFI) associated with CD4 serves as a surrogate for CD4 surface expression. Medians for each condition are presented. Statistical significance was calculated by paired comparisons between each treated condition (Wilcoxon test).

**Figure S8:**
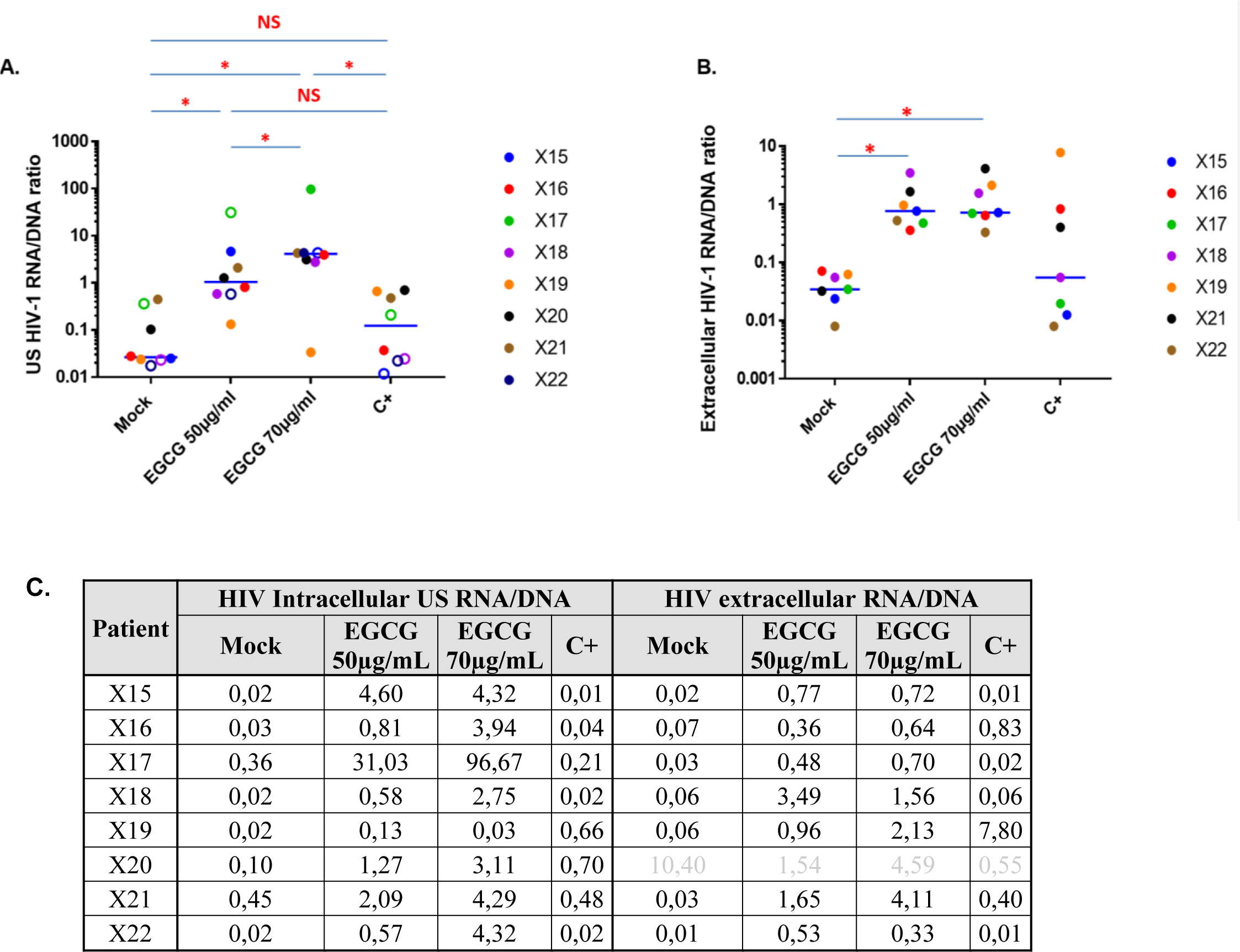
HIV-1 proviruses are statistically more transcriptionally active in EGCG-treated *ex vivo* patient cell cultures. **(A)** Comparison of HIV-1 intracellular US RNA/DNA ratios between mock-treated, EGCG 50µg/mL, EGCG 70µg/mL and positive control stimulations of *ex vivo* patient X15 to X22 cultures is presented. **(B)** Comparison of HIV-1 extracellular RNA/DNA ratios between mock-treated, EGCG 50µg/mL, EGCG 70µg/mL and positive control stimulations of *ex vivo* patient X15 to X22 cultures is presented. In both panels, statistical significance was calculated by paired comparisons between each treated condition (Wilcoxon test). **(C)** The ratios of HIV intracellular US RNA/DNA or extracellular RNA/DNA were calculated for the individuals X15 to X22. Of note, individual X20 was excluded due to high level of extracellular HIV-1 RNA in mock-treated condition at 24h post-stimulation.

**Figure S9:**
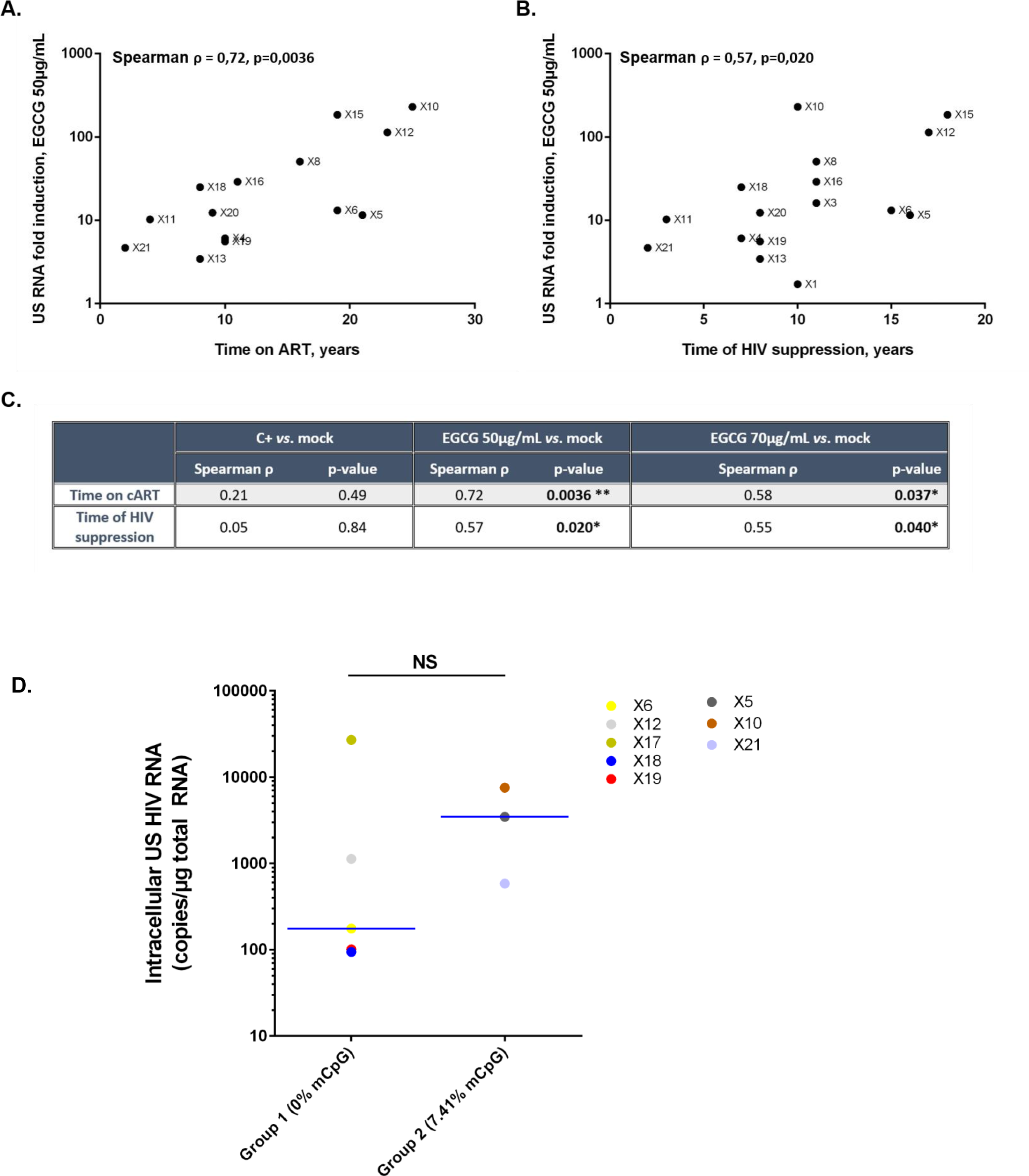
EGCG reactivation potency *ex vivo* correlates to temporal parameters of infection in HIV^+^ individuals and follows the DNA methylation level of the viral promoter. For HIV^+^ individuals presenting detectable values of cell-associated HIV-1 US RNA, fold inductions over mock condition were calculated, based on the data presented in Fig. 5A. Spearman correlations between quantity of cell-associated HIV-1 US RNA for 50µg/mL of EGCG and time of treatment **(A)** or time as aviremic **(B)** were calculated based on the data presented in Table S1. In **(C)**, raw values of Spearman ρ and associate p-values for each condition are presented. **(D)**The methylation status for 8 individuals out of 22 was obtained by sodium bisulfite sequencing. We clustered individuals in groups presenting non-methylated or methylated 5’LTR and analyzed their respective median EGCG reactivation capacity, in terms of cell-associated HIV-1 US RNA for 50µg/mL of EGCG.

**Table S1:**
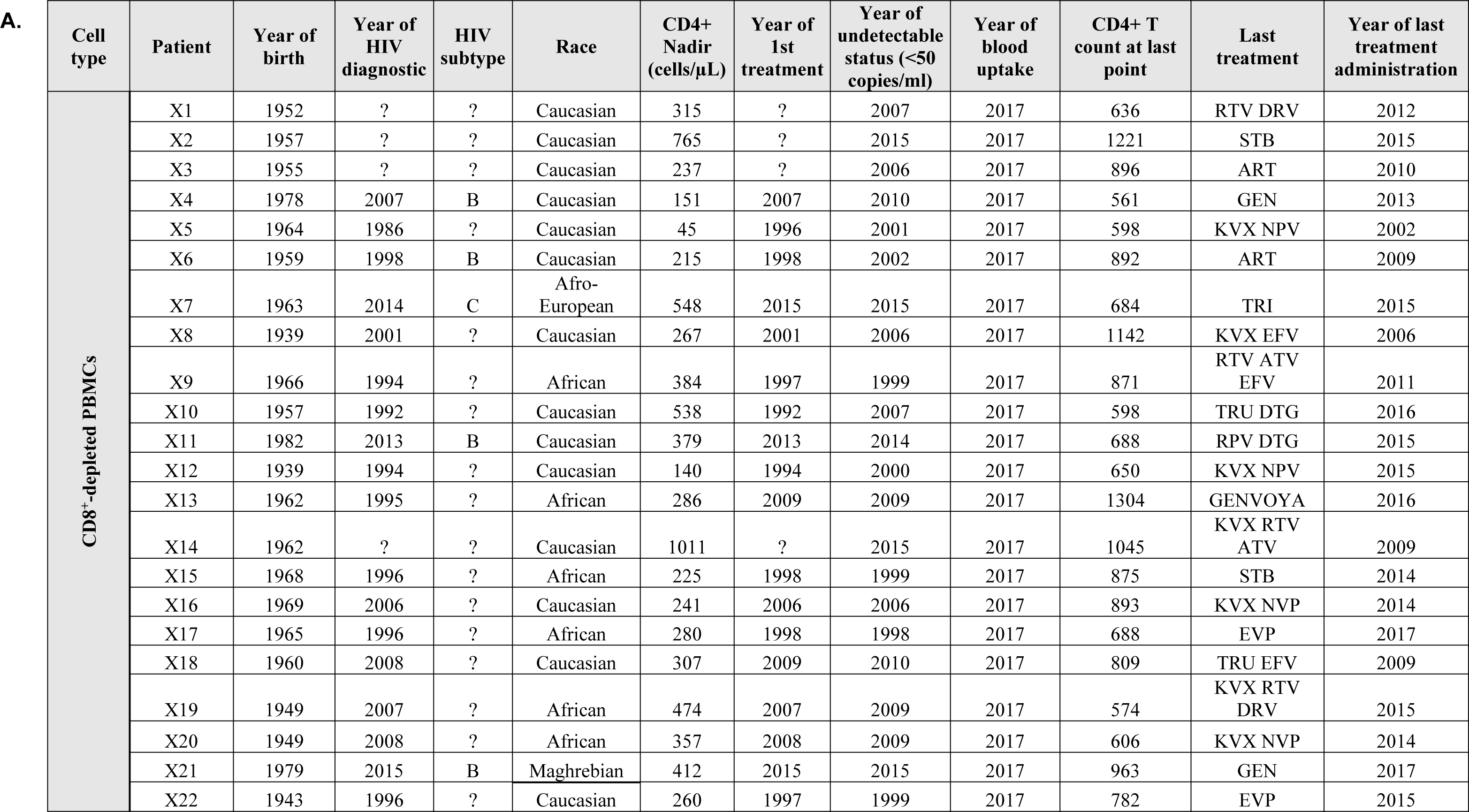
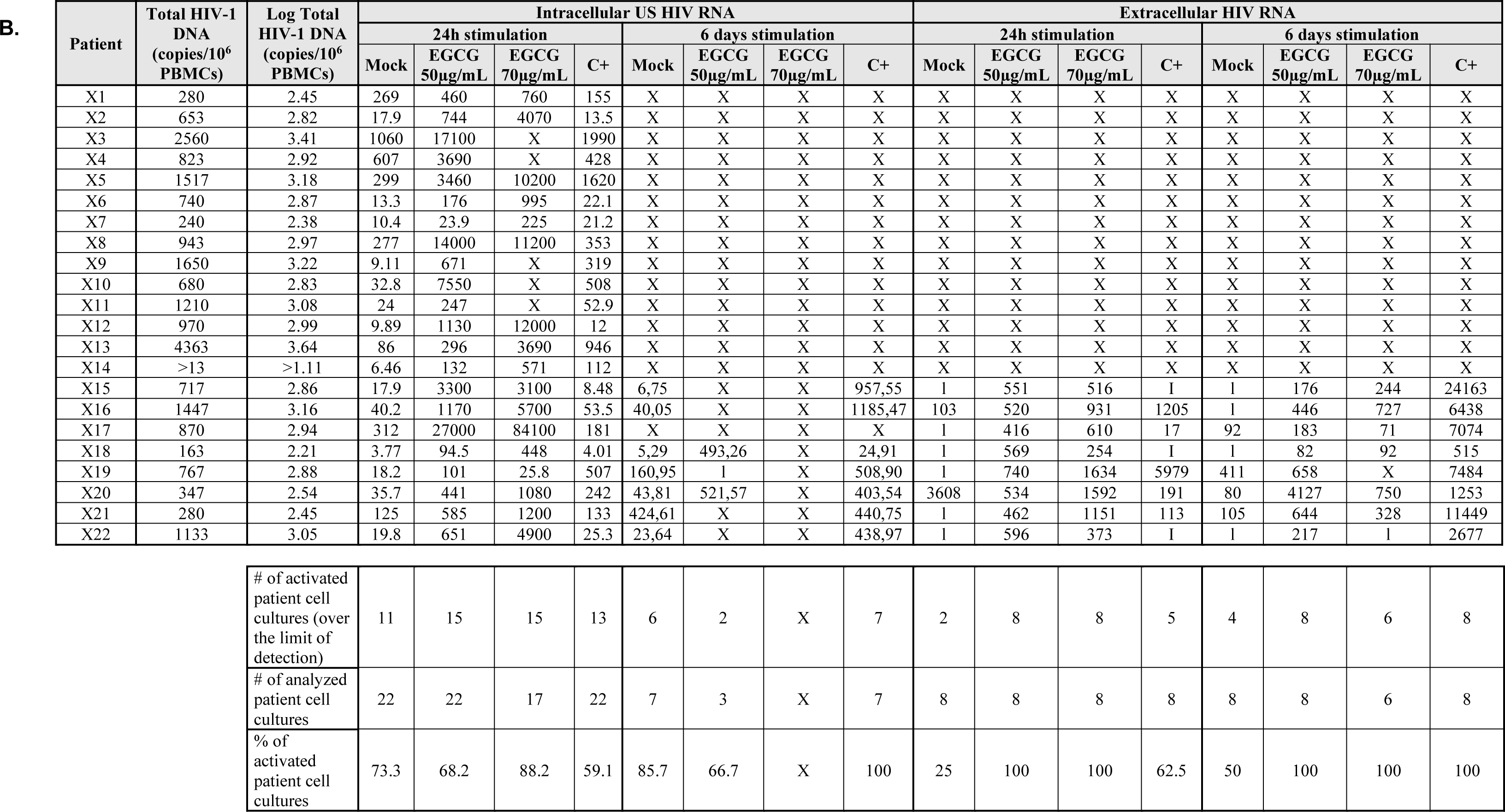
EGCG induces HIV-1 recovery in CD8^+^-depleted PBMCs from aviremic cART-treated HIV^+^ individuals. **(A)** HIV+ individuals’ clinical characteristics are listed. ART= antiretroviral therapy, ATV= atazanavir, DRV= darunavir, DTG= dolutegravir EFV= efavirenz, EVP= eviplera (rilpivirine, emtricitabine, tenofovir), GENVOYA= elvitegravir/cobicistat/emtricitabine/tenofovir, KVX = kivexa, NPV = nevirapine, RTV = ritonavir, RPV=rilpivirine, STB = elvitegravir, cobicistat, emtricitabine, tenofovir, TRI= triple cocktail, TRU = Truvada and “?” = unknown. **(B)** *Ex vivo* cultures of CD8^+^-depleted PBMCs purified from blood of 22 aviremic cART-treated HIV^+^ individuals were mock-treated or treated with anti-CD3+anti-CD28 antibodies, as positive control stimulation, or with EGCG 50µg/mL or EGCG 70µg/mL. Twenty-four hours or six days post-treatment, concentrations of intracellular unspliced HIV- 1 RNA or of extracellular HIV-1 RNA was determined, respectively in cells and in cell culture supernatants. Total HIV-1 DNA was expressed as HIV-1 DNA copies *per* million CD8^+^-depleted PBMCs. ‘I’ indicates that the value is below the limit of detection. ‘X’ indicates that the condition was not tested. Of note, individual X20 was excluded from extracellular quantification due to high level of extracellular HIV-1 RNA in mock-treated condition at 24h post-stimulation.

| Cell type | Patient | Year of birth | Year of HIV diagnostic | HIV subtype | Race | CD4+ Nadir (cells/μL) | Year of 1st treatment | Year of undetectable status (<50 copies/ml) | Year of blood uptake | CD4+ T count at last point | Last treatment | Year of last treatment administration |
| --- | --- | --- | --- | --- | --- | --- | --- | --- | --- | --- | --- | --- |
| CD8<sup>+</sup>-depleted PBMCs | X1 | 1952 | ? | ? | Caucasian | 315 | ? | 2007 | 2017 | 636 | RTV DRV | 2012 |
|  | X2 | 1957 | ? | ? | Caucasian | 765 | ? | 2015 | 2017 | 1221 | STB | 2015 |
|  | X3 | 1955 | ? | ? | Caucasian | 237 | ? | 2006 | 2017 | 896 | ART | 2010 |
|  | X4 | 1978 | 2007 | B | Caucasian | 151 | 2007 | 2010 | 2017 | 561 | GEN | 2013 |
|  | X5 | 1964 | 1986 | ? | Caucasian | 45 | 1996 | 2001 | 2017 | 598 | KVX NPV | 2002 |
|  | X6 | 1959 | 1998 | B | Caucasian | 215 | 1998 | 2002 | 2017 | 892 | ART | 2009 |
|  | X7 | 1963 | 2014 | C | Afro-European | 548 | 2015 | 2015 | 2017 | 684 | TRI | 2015 |
|  | X8 | 1939 | 2001 | ? | Caucasian | 267 | 2001 | 2006 | 2017 | 1142 | KVX EFV | 2006 |
|  | X9 | 1966 | 1994 | ? | African | 384 | 1997 | 1999 | 2017 | 871 | RTV ATV EFV | 2011 |
|  | X10 | 1957 | 1992 | ? | Caucasian | 538 | 1992 | 2007 | 2017 | 598 | TRU DTG | 2016 |
|  | X11 | 1982 | 2013 | B | Caucasian | 379 | 2013 | 2014 | 2017 | 688 | RPV DTG | 2015 |
|  | X12 | 1939 | 1994 | ? | Caucasian | 140 | 1994 | 2000 | 2017 | 650 | KVX NPV | 2015 |
|  | X13 | 1962 | 1995 | ? | African | 286 | 2009 | 2009 | 2017 | 1304 | GENVOYA | 2016 |
|  | X14 | 1962 | ? | ? | Caucasian | 1011 | ? | 2015 | 2017 | 1045 | KVX RTV ATV | 2009 |
|  | X15 | 1968 | 1996 | ? | African | 225 | 1998 | 1999 | 2017 | 875 | STB | 2014 |
|  | X16 | 1969 | 2006 | ? | Caucasian | 241 | 2006 | 2006 | 2017 | 893 | KVX NVP | 2014 |
|  | X17 | 1965 | 1996 | ? | African | 280 | 1998 | 1998 | 2017 | 688 | EVP | 2017 |
|  | X18 | 1960 | 2008 | ? | Caucasian | 307 | 2009 | 2010 | 2017 | 809 | TRU EFV | 2009 |
|  | X19 | 1949 | 2007 | ? | African | 474 | 2007 | 2009 | 2017 | 574 | KVX RTV DRV | 2015 |
|  | X20 | 1949 | 2008 | ? | African | 357 | 2008 | 2009 | 2017 | 606 | KVX NVP | 2014 |
|  | X21 | 1979 | 2015 | B | Maghrebian | 412 | 2015 | 2015 | 2017 | 963 | GEN | 2017 |
|  | X22 | 1943 | 1996 | ? | Caucasian | 260 | 1997 | 1999 | 2017 | 782 | EVP | 2015 |

B.
| Patient | Total HIV-1 DNA<br>(copies/10<sup>6</sup> PBMCs) | Log Total HIV-1 DNA<br>(copies/10<sup>6</sup> PBMCs) | Intracellular US HIV RNA |  |  |  |  |  |  |  | Extracellular HIV RNA |  |  |  |  |  |  |  |
| --- | --- | --- | --- | --- | --- | --- | --- | --- | --- | --- | --- | --- | --- | --- | --- | --- | --- | --- |
|  |  |  | 24h stimulation |  |  |  | 6 days stimulation |  |  |  | 24h stimulation |  |  |  | 6 days stimulation |  |  |  |
|  |  |  | Mock | EGCG<br>50µg/mL | EGCG<br>70µg/mL | C+ | Mock | EGCG<br>50µg/mL | EGCG<br>70µg/mL | C+ | Mock | EGCG<br>50µg/mL | EGCG<br>70µg/mL | C+ | Mock | EGCG<br>50µg/mL | EGCG<br>70µg/mL | C+ |
| X1 | 280 | 2.45 | 269 | 460 | 760 | 155 | X | X | X | X | X | X | X | X | X | X | X | X |
| X2 | 653 | 2.82 | 17.9 | 744 | 4070 | 13.5 | X | X | X | X | X | X | X | X | X | X | X | X |
| X3 | 2560 | 3.41 | 1060 | 17100 | X | 1990 | X | X | X | X | X | X | X | X | X | X | X | X |
| X4 | 823 | 2.92 | 607 | 3690 | X | 428 | X | X | X | X | X | X | X | X | X | X | X | X |
| X5 | 1517 | 3.18 | 299 | 3460 | 10200 | 1620 | X | X | X | X | X | X | X | X | X | X | X | X |
| X6 | 740 | 2.87 | 13.3 | 176 | 995 | 22.1 | X | X | X | X | X | X | X | X | X | X | X | X |
| X7 | 240 | 2.38 | 10.4 | 23.9 | 225 | 21.2 | X | X | X | X | X | X | X | X | X | X | X | X |
| X8 | 943 | 2.97 | 277 | 14000 | 11200 | 353 | X | X | X | X | X | X | X | X | X | X | X | X |
| X9 | 1650 | 3.22 | 9.11 | 671 | X | 319 | X | X | X | X | X | X | X | X | X | X | X | X |
| X10 | 680 | 2.83 | 32.8 | 7550 | X | 508 | X | X | X | X | X | X | X | X | X | X | X | X |
| X11 | 1210 | 3.08 | 24 | 247 | X | 52.9 | X | X | X | X | X | X | X | X | X | X | X | X |
| X12 | 970 | 2.99 | 9.89 | 1130 | 12000 | 12 | X | X | X | X | X | X | X | X | X | X | X | X |
| X13 | 4363 | 3.64 | 86 | 296 | 3690 | 946 | X | X | X | X | X | X | X | X | X | X | X | X |
| X14 | >13 | >1.11 | 6.46 | 132 | 571 | 112 | X | X | X | X | X | X | X | X | X | X | X | X |
| X15 | 717 | 2.86 | 17.9 | 3300 | 3100 | 8.48 | 6.75 | X | X | 957,55 | 1 | 551 | 516 | 1 | 1 | 176 | 244 | 24163 |
| X16 | 1447 | 3.16 | 40.2 | 1170 | 5700 | 53.5 | 40,05 | X | X | 1185,47 | 103 | 520 | 931 | 1205 | 1 | 446 | 727 | 6438 |
| X17 | 870 | 2.94 | 312 | 27000 | 84100 | 181 | X | X | X | X | 1 | 416 | 610 | 17 | 92 | 183 | 71 | 7074 |
| X18 | 163 | 2.21 | 3.77 | 94.5 | 448 | 4.01 | 5,29 | 493,26 | X | 24,91 | 1 | 569 | 254 | 1 | 1 | 82 | 92 | 515 |
| X19 | 767 | 2.88 | 18.2 | 101 | 25.8 | 507 | 160,95 | 1 | X | 508,90 | 1 | 740 | 1634 | 5979 | 411 | 658 | X | 7484 |
| X20 | 347 | 2.54 | 35.7 | 441 | 1080 | 242 | 43,81 | 521,57 | X | 403,54 | 3608 | 534 | 1592 | 191 | 80 | 4127 | 750 | 1253 |
| X21 | 280 | 2.45 | 125 | 585 | 1200 | 133 | 424,61 | X | X | 440,75 | 1 | 462 | 1151 | 113 | 105 | 644 | 328 | 11449 |
| X22 | 1133 | 3.05 | 19.8 | 651 | 4900 | 25.3 | 23,64 | X | X | 438,97 | 1 | 596 | 373 | 1 | 1 | 217 | 1 | 2677 |

|  |  |  |  |  |  |  |  |  |  |  |  |  |  |  |  |  |
| --- | --- | --- | --- | --- | --- | --- | --- | --- | --- | --- | --- | --- | --- | --- | --- | --- |
| # of activated patient cell cultures (over the limit of detection) | 11 | 15 | 15 | 13 | 6 | 2 | X | 7 | 2 | 8 | 8 | 5 | 4 | 8 | 6 | 8 |
| # of analyzed patient cell cultures | 22 | 22 | 17 | 22 | 7 | 3 | X | 7 | 8 | 8 | 8 | 8 | 8 | 8 | 6 | 8 |
| % of activated patient cell cultures | 73.3 | 68.2 | 88.2 | 59.1 | 85.7 | 66.7 | X | 100 | 25 | 100 | 100 | 62.5 | 50 | 100 | 100 | 100 |

